# High-fidelity generalizable light-field reconstruction of biological dynamics with physics-informed meta neural representation

**DOI:** 10.1101/2023.11.25.568636

**Authors:** Chengqiang Yi, Jiahao Sun, Minglu Sun, Lanxin Zhu, Yifan Ma, Sicen Wu, Yuting Liu, Shangbang Gao, Meng Zhang, Yuhui Zhang, Zhaoqiang Wang, Hsiai K. Tzung, Peng Zou, Binbing Liu, Dongyu Li, Peng Fei

## Abstract

Light-field microscopy (LFM) captures 3D biological dynamics by single 2D snapshots but suffers from limited resolution and artifacts during 2D-to-3D inversion. Here, we introduce light-field meta neural representation (LFMNR), a novel self-supervised paradigm that utilizes physics-informed light-field implicit neural representation (LFINR) and meta learning for high-quality 3D reconstruction in Fourier-LFM. By developing a physics-based hybrid-rendering model, LFINR achieves artifact-free light-field reconstruction with enhanced spatial resolution (>1.4-fold improvement). Additionally, the integration of meta-learning and progressive sampling strategies mitigates INR’s intrinsic limitations in low reconstruction speed caused by scene-specific optimization, enabling a ∼100-fold acceleration in the representation of consecutive volumes and facilitating the visualization of sustained 3D dynamics. These advancements enable LFMNR to deliver superior imaging with high spatiotemporal resolution and low phototoxicity, as demonstrated by capturing instantaneous voltage signals in *C. elegans* at 100 volumes per second and recording 6000 time points of organelle dynamics over 25 hours.

## Introduction

Understanding the spatial dynamics and interactions within microenvironments necessitates high spatiotemporal resolution across three dimensions. This imperative drives the advancement of volumetric imaging technologies to probe rapid biological processes, such as cardiac hemodynamics^1, 2^ and intracellular interactions^3–5^. While scanning-based microscopy compromises volumetric imaging speed and can induce additional phototoxicity due to its sequential scanning approach, light field microscopy^6^ (LFM) enables one-shot volumetric imaging by capturing all fluorescence signals from an excited volume in a single 2D image through multi-view captures. Though LFM is ideal for imaging dynamic biological processes^7^, the non-uniform spatial resolution resulting from frequency aliasing limits its broader application. Fourier light field microscopy^8–10^ (FLFM) employs a microlens array at the Fourier plane to mitigate spatial sampling aliasing in standard LFM and better leverage the optical aperture, thereby achieving improved spatial resolution with fewer views. Since these methods commonly employ Richard-Lucy (RL) deconvolution for iterative 3D volume reconstruction using the point spread function, challenges arise under the lower angular sampling rate and limited angle-range of FLFM. Such iterative-based algorithms, relying solely on imaging model priors for limited-view 3D reconstruction, tend to introduce artifacts and compromise structure fidelity^11–14^.

To address limitations in multi-view 3D reconstruction, data-driven algorithms are proposed, leveraging deep learning models to convert 2D LF captures into high-resolution 3D stacks with improved reconstruction quality^15–19^. However, end-to-end supervised approaches require large-scale volumetric data and well-aligned “2D-3D” training pairs, which are not readily accessible and prone to introducing artifacts due to mismatch between training and experimental data^20^. Self-supervised learning-based algorithms, which do not require volumetric data, offer a promising alternative for advanced light-field reconstruction. Current self-supervised methods in microscopy tasks often integrate physical priors into models through well-designed loss functions derived from analytical models to ensure physical consistency^21–23^. However, building self-supervised models for 3D reconstruction encounters challenges primarily due to the dimensionality reduction, which adds complexity to finding viable solutions within physical consistency constraints. This complexity often leads to significant artifacts, hallucinations, and discontinuities in the reconstructed output (**Supplementary** Fig. 1, **Supplementary Note 1**).

Recently, Neural Radiance Field^24^ (NeRF) has been known for its ability to generate novel views with geometry consistency, and applied in various vision tasks^25–28^. NeRF employs a Multi-Layer Perceptron (MLP) to represent 3D scenes from 2D multi-view input, known as implicit neural representation (INR), capturing the latent continuous function of signal distribution. While NeRF’s volumetric representation correlates with the 3D reconstruction inverse problem in FLFM, differences in forward models between macrophotography and diffraction theory in microscopy imaging pose big challenges for realization of INR-based light-field reconstruction. Moreover, NeRF’s grid-based training strategy is highly computationally intensive, limiting its further application in processing dynamic data.

Here, we introduce a physically informed, meta-learning based self-supervised approach, termed light-field meta neural representation (LFMNR), for efficient Fourier light-field 3D reconstruction with high fidelity. First, we developed a physics-based hybrid-rendering model as an optimization constraint to achieve light field reconstruction with implicit neural representation (LFINR) for the first time. Leveraging the continuity of the INR function and the geometry consistency of synthesized views, LFINR effectively reduces artifacts and axial elongation even with extremely-limited views input of 3, far surpassing classical model-based light-field deconvolution approaches with approximately >1.3-fold structural fidelity improvement and >1.4-fold resolution enhancement. Then, we employ a progressive sampling strategy with structural error maps to significantly reduce computational time compared to conventional INR methods. Additionally, based on the intrinsic continuity of dynamic LF time-sequences, we utilize meta-learning to transfer optimized weights to adjacent time points. The combination of progressive sampling and meta learning enables ∼100x accelerated light-field meta neural representation (LFMNR) for high-throughput reconstruction of sustained light-field videos. Long-term cell apoptosis and various intracellular dynamic events, including mitochondrial fission, mitochondria-lysosome interactions, and peroxisome dynamics, have been identified and quantitatively analyzed to showcase the capabilities of LFMNR. Furthermore, the self-supervised LFMNR also enables high-resolution reconstruction of instantaneous 3D voltage signaling in the pharyngeal muscles of *C. elegans* during pumping behavior with millisecond temporal resolution.

## Results

### The principle of LFINR

The framework of LFINR includes 3 steps: spatial coordinates sampling, INR network construction, and physics-model consistency evaluation, as illustrated in **Fig. 1** and **Supplementary** Fig. 2. Unlike conventional convolution neural networks (CNN) in image enhancement or reconstruction, LFINR requires spatial coordinates }*^N^_i=1_* as network input, which means a 3D spatial grid covering the reconstructed signals needed to be defined before network inference (“*Objective space*” in **Fig. 1a**). However, due to the discrete grid form of image data, the quantity of spatial coordinates will increase multiplicatively with the enlargement of image size, which burden the computation cost and reduce network convergence speed. Here, we introduced a progressive sampling strategy, which first used down-sampled mesh grid to pre-train network and next fed all grids for details fitting, to make network fast convergence. Initially, a sparse-sampled mesh grid was employed to pre-train the network, providing a preliminary approximation of the overall distribution (“*i. Sparse sampling*” in **Fig. 1a**). Subsequently, during the fine-tuning phase, the full-size grids were firstly fed into the network for details recovery (“*ii. Full sampling*” in **Fig. 1a**).

**Figure 1.**
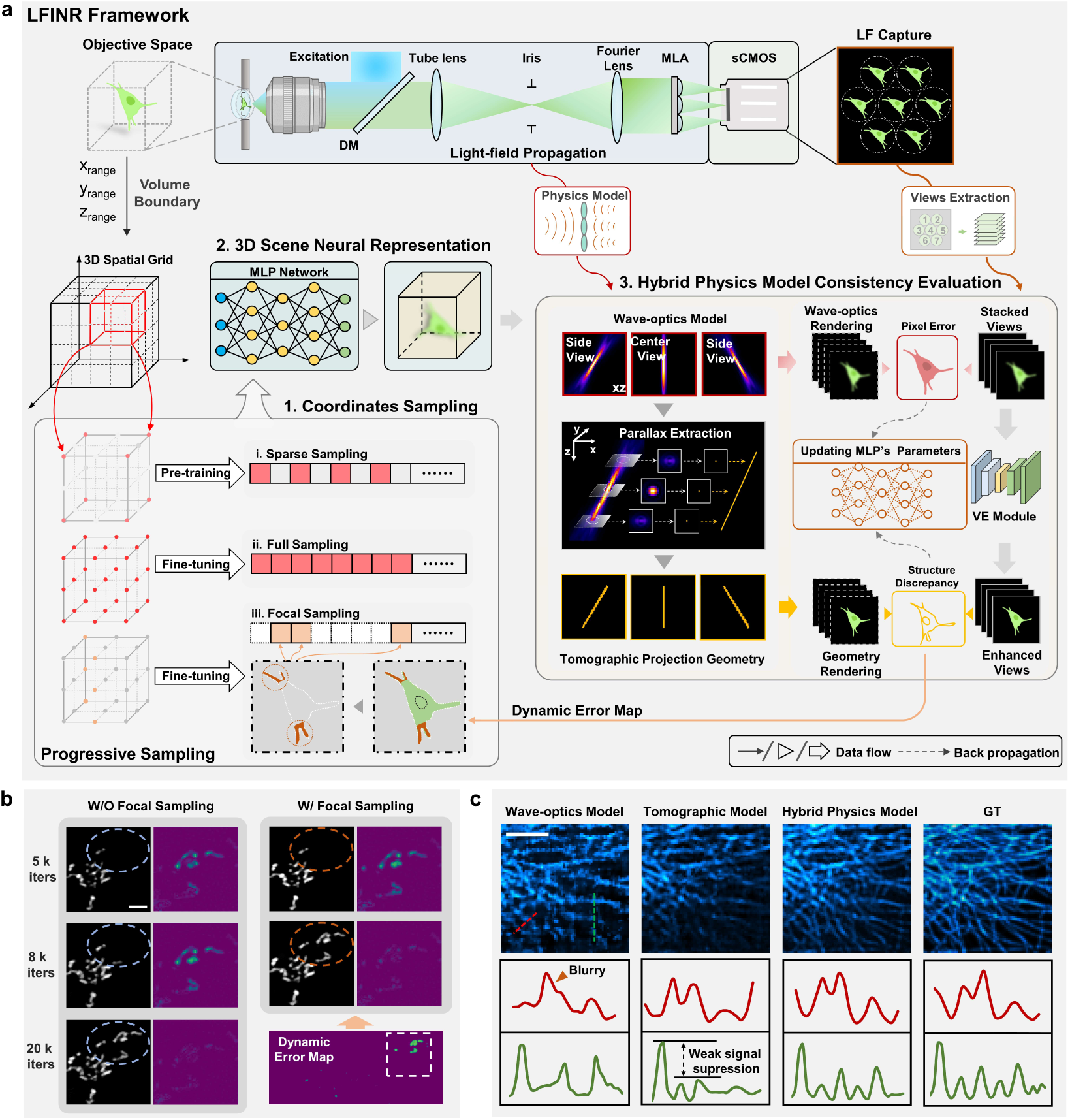
The overview of LFINR framework. a, Workflow of LFINR, which contains 3 steps: Sampling coordinates from 3D spatial grid with progressive strategy; Constructing an MLP based scene-representation network; Computing the hybrid physics model consistency. With constant network optimization in a self-supervised manner, LFINR could learn to represent the 3D scene within the MLP, referred to implicit neural representation. During the optimization process, progressive sampling strategy was adopted to accelerate network convergence under the guidance of structural error map. The hybrid physics model contains wave-optics model for evaluating pixel errors between the extracted views and wave-optics rendering results, and tomographic projection geometry for evaluating structure discrepancy between geometry rendering results and enhanced views obtained by view-enhancement (VE) module. **b**, Focal sampling strategy enhances signal integrity and accelerates convergence in 3D reconstruction of mitochondria. Scale bar: 5 μm. **c**, Maximum intensity projections (MIPs) of microtubule reconstructed via wave-optics model based INR, tomographic model based INR and hybrid physics model based INR (LFINR). Scale bar: 5 μm.

Then, during the later stages of training, a focal sampling approach was implemented to prioritize the training on regions exhibiting higher distortion, in which, we leveraged local structural similarity to compute the error map (“*iii. Focal sampling*” in **Fig. 1a, Methods**). With this spatially aware optimization strategy, LFINR could achieve ∼2-fold acceleration and outperform in weak-signal reconstruction when compared with random-sampling based approach^24^ (**Fig. 1b, Supplementary** Fig. 3). After coordinates sampling, a fully-connected network (“*MLP network*” in **Fig. 1a**) was constructed to serve as the underlying continuous volumetric scene function. By querying the 3D spatial coordinates, the network could recover the volumetric intensity information.

Finally, the produced volume was utilized to generate projections with forward rendering model for physical consistency evaluation. However, during this process, the difference between forward rendering model and real-world model notably degraded the 3D reconstruction. For example, the photography model utilized in conventional NeRF^24^ relies on ray-casting model to form the volumetric density and color of opaque 3D scene. While FLFM imaging model is based on wave-optics theory and aims to accurately describe the intensity distribution of semi-/transparent sample. The ignoration of diffraction effect in wave-optics model will blur the represented 3D volume with increased errors (**Supplementary** Fig. 4).

Taking this into account, we developed a hybrid physics model following the principle of wave-optics model and tomographic projection geometry to obtain dual-channel forward projections in rendering process and compute the hybrid loss function for evaluating physics consistency **(Fig. 1a, Methods**). For the wave-optics rendering branch, the FLFM Point Spread Function (PSF) was employed to convolve with the predicted volume to generate wave-optics rendering results, which simulated the light-field propagation determined by the hardware settings (“*Light-field propagation*”, **Fig. 1a**). For the geometry rendering branch, we derived the projection matrix through parallax extraction and obtained the geometry rendering results (**Methods**). During the model optimization phase, we adopted weighted hybrid function based on the dual rendering results. The first term of loss function was pixel-wise errors (L2 loss) between the extracted views and wave-optics rendering results. The second one was structure discrepancy to capture the skeleton information of input views. Before this loss computation, we employed a self-supervised view-enhancement network, termed as VE module in **Fig. 1a**, to remove the depth-correlated blur in extracted LF views, which matched the geometry rendering model (**Supplementary Note 2**). Then, the structure discrepancy was derived from structure similarity index measurement (SSIM) between enhanced views and geometry rendering results. By continuously minimizing the hybrid loss, LFINR gradually learned to produce the 3D volume only with input coordinates, in a form of scene representation. Our hybrid physics projection model leverages the complementary strengths of the dual branches while mitigating their respective weaknesses—blurring in wave-optics model results and weak signal loss in tomographic model results (**Fig. 1b**). This synergy improves reconstruction quality and accelerates network convergence (**Supplementary** Fig. 5, **Supplementary Note 3**). Further details about the network implementations can be found in the **Methods** section.

### LFINR enables high-fidelity 3D reconstruction by its continuous representation

For the inverse problem of 3D reconstruction from encoded multi-view light-field images, the quality of reconstructed results is highly depended on the performance of decoded algorithm. The 3D supervised DL algorithms^15, 17, 19^ solve such problem in an unexplainable way by building CNN network mapping from 2D capture to 3D volume, which suffers the generalization errors due to the data bias^20^. The classical model based method, such as RL-deconvolution^8^, is still the most widely-used approach especially when the volumetric data guidance is not available. However, the inherent constraints within inverse problems, characterized by limited angle range and sparse sampling of captured views^29^, often lead to poor reconstruction quality, particularly evident along the axial dimension and for densely packed signals.

To assess the performance of various approaches, we compared the reconstruction results on different biological samples using numerical simulations (**Fig. 2**). As shown in the simulation workflow (**Fig. 2a**), 3D ground truth (GT) data were obtained using a high-resolution scanning-based microscope (**Methods**) and used to generate Fourier light-field images under different FLFM setups. These simulated light-field images were subsequently reconstructed into volumes through various 3D reconstruction algorithms, and the accuracy of these reconstructions was evaluated by comparison with the GT. In **Fig. 2b**, LFINR preserved obviously more structure details and higher contrast compared with FLFM deconvolution (**Methods**), abbreviated as Deconv, when decoding the 3D microtubule signals in an U2OS cells (labeled with 3mEmerald-Ensconsin) from simulated 7 FLFM views (**Methods**). The Deconv results showed noticeable blurs in lateral and axial projections, along with significant view-related stripe artifacts. These issues were confirmed by the narrow bandwidth in the Fourier spectrum images and the low SSIM values (“*Deconv*” in **Fig. 2c**). Such deficiencies came from the depth-wise blurry modulation and signal cross-talk in Fourier light-field imaging process. To suppress the blurring effect, we employed a self-supervised view-enhancement (VE) network to remove the resolution degradation in FLFM views (**Supplementary Note 2**). The deblurred views were subsequently deconvolved using the geometric projection matrix, termed VE-Deconv, to generate 3D reconstructions (**Methods**). The blur occurred in Deconv results decreased remarkably but the striped artifacts still existed as shown in “*VE-Deconv*”, **Fig. 2c**. In sharp contrast, LFINR achieved artifacts-free, high-fidelity 3D reconstruction, with a higher SSIM of ∼0.97 and a lower Normalized Root Mean Square Error (NRMSE) of ∼0.056. It also recovered substantially more high-frequency structural details, which also further demonstrated that LFINR offers superior optical sectioning capability compared to deconvolution-based approaches (**Supplementary** Fig. 6). These notable quality improvements stemmed from the intrinsic continuity of implicit representation and physics-based hybrid rendering model in LFINR (**Supplementary** Fig. 7). Furthermore, we also compared LFINR with VCD^15^, a state-of-the-art deep-learning approach for light-field reconstruction based on volume-supervised learning (**Fig. 2c, Supplementary** Fig. 1). The results revealed that, while both methods demonstrated similar lateral structural fidelity, VCD reconstructions showed reduced SSIM and increased NRMSE values in axial planes. This decline in performance was ascribed to the inherent data bias in the volume-supervised approach, a limitation that has been circumvented in our self-supervised LFINR.

**Figure 2.**
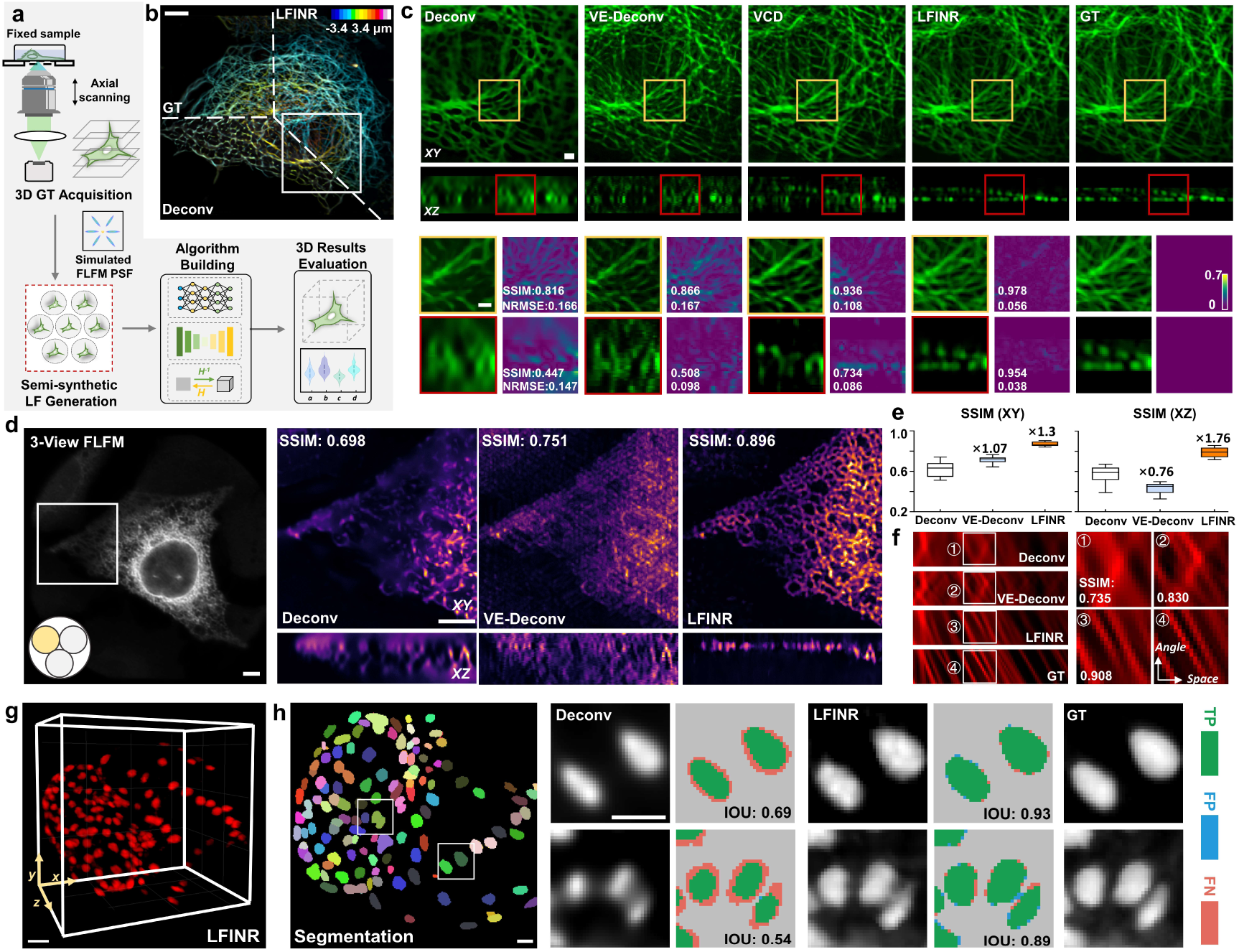
Light-field reconstruction with improve quality by LFINR. a, The workflow for comparing 3D reconstruction algorithms using numerical simulations consists of the following steps: (i) acquiring 3D ground truth (GT), (ii) generating semi-synthetic light-field images, (iii) reconstructing 3D signals using various algorithms, and (iv) evaluating reconstruction quality. **b,** The depth-encoded projections of microtubules by LFINR reconstruction, FLFM deconvolution (Deconv) and ground truth. The number of FLFM views used was 7. **c,** Magnified horizontal and vertical sectional views of white rectangular boxes marked in (**b)**, denoting the higher structure fidelity of LFINR results compared to Deconv, view-enhanced deconvolution (VE-Deconv) and VCD results. **d,** The comparison between Deconv, VE-Deconv and LFINR results reconstructed from three FLFM views of endoplasmic reticulum (ER). **e,** Structure similarity comparison of three approaches. The numbers above the “VE-Deconv” and “LFINR” boxes represent the structure similarity enhancement ratio compared to the Deconv results. **f,** Angular consistency evaluation via comparison of computed epi-polar plane images (EPIs) across different approaches. Top to bottom: Deconv, VE-Deconv, LFINR and GT. **g,** LFINR reconstruction from 19 FLFM views of cardiomyocyte nuclei**. h,** The segmentation results of reconstructed volume in **(g)**, along with the comparison of Intersection over Union (IOU) scores between Deconv and LFINR. TP: True Positive. FP: False Positive. FN: False Negative. Scale bars: 50 μm in (**b**), 2 μm in (**c**), 5 μm in (**d**), 20 μm in (**g**), 10 μm in (**h**).

In addition to the improvements in structural fidelity and spatial resolution, LFINR, with its ability for arbitrary view synthesis facilitated by the continuity of representation, also demonstrate significant superiority in 3D reconstruction of ER (labeled with EGFP-Sec61β) with extremely low number of light-field views, such as just 3, achieving ∼2.3-fold higher SSIM, ∼1.3-fold improvement in xy planes and ∼1.76-fold improvement in xz planes (**Figs. 2d-e, Supplementary** Fig. 8). The deconvolution-based results exhibited severe striped artifacts in axial plane (white arrowheads), when only 3 views were used. Such undesirable hallucinations across different depth further deteriorated the lateral structural fidelity denoted by the lower SSIM (**Figs. 2e**). In addition to space domain evaluation, we compared the geometry consistency of epi-polar plane images (EPIs)^30^ generated by different approaches (**Fig. 2f, Methods, Supplementary** Fig. 9). These EPIs represent the distribution of both angle and space, with the slopes of the lines indicating parallax across different views. When angular sampling was reduced, deconvolution-based methods exhibited artificial stripes in EPIs. In contrast, LFINR’s outputs maintained a seamless transition across various views, as evidenced by the consistent alignment between GT’s EPIs and LFINR one denoted by a higher SSIM of 0.908. This highlight LFINR’s ability to generate arbitrary views with strong geometric consistency, even under the constraints of limited view supervision

We noted that, with the image quality enhancement, LFINR also achieves higher accuracy in downstream analysis (**Figs. 2g, h**). The segmentation results of cardiomyocyte nuclei (labeled with *Tg (myl7:nls-gfp)*) showed higher intersection over union (IOU) scores of LFINR, while Deconv showed obvious errors in the boundary of single cell (red area in “*Deconv*”, **Fig. 2g**), indicating the structural errors in Deconv.

It is noteworthy that beyond the application on FLFM, we also validated the self-supervised INR approach across various 3D computational imaging modalities, such as optical projection tomography (OPT)^31^, Fourier ptychographic microscopy (FPM)^32, 33^ and cryo-electron tomography (Cryo-ET)^34–36^. The results consistently demonstrated superior structural fidelity compared to conventional model-based approaches, highlighting the versality of INR in tackling inverse tomographic reconstruction problems (**Supplementary** Figs. 10-13).

### Spatial-angular bandwidth enlargement offered by LFINR

The hardware design of Fourier light-field microscopy (FLFM) is an art of balance on angular and spatial information in a limited bandwidth determined by objective lens. In the image domain, such trade-off usually reflected as the form of spatial resolution degradation and axial artifacts. Here, in a complete self-supervised way eliminating concerns regarding generalization, LFINR showcased its strong reconstruction capability across diverse FLFM imaging configurations. The superior results were quantitatively verified through various metrics (**Fig. 3**).

**Figure 3.**
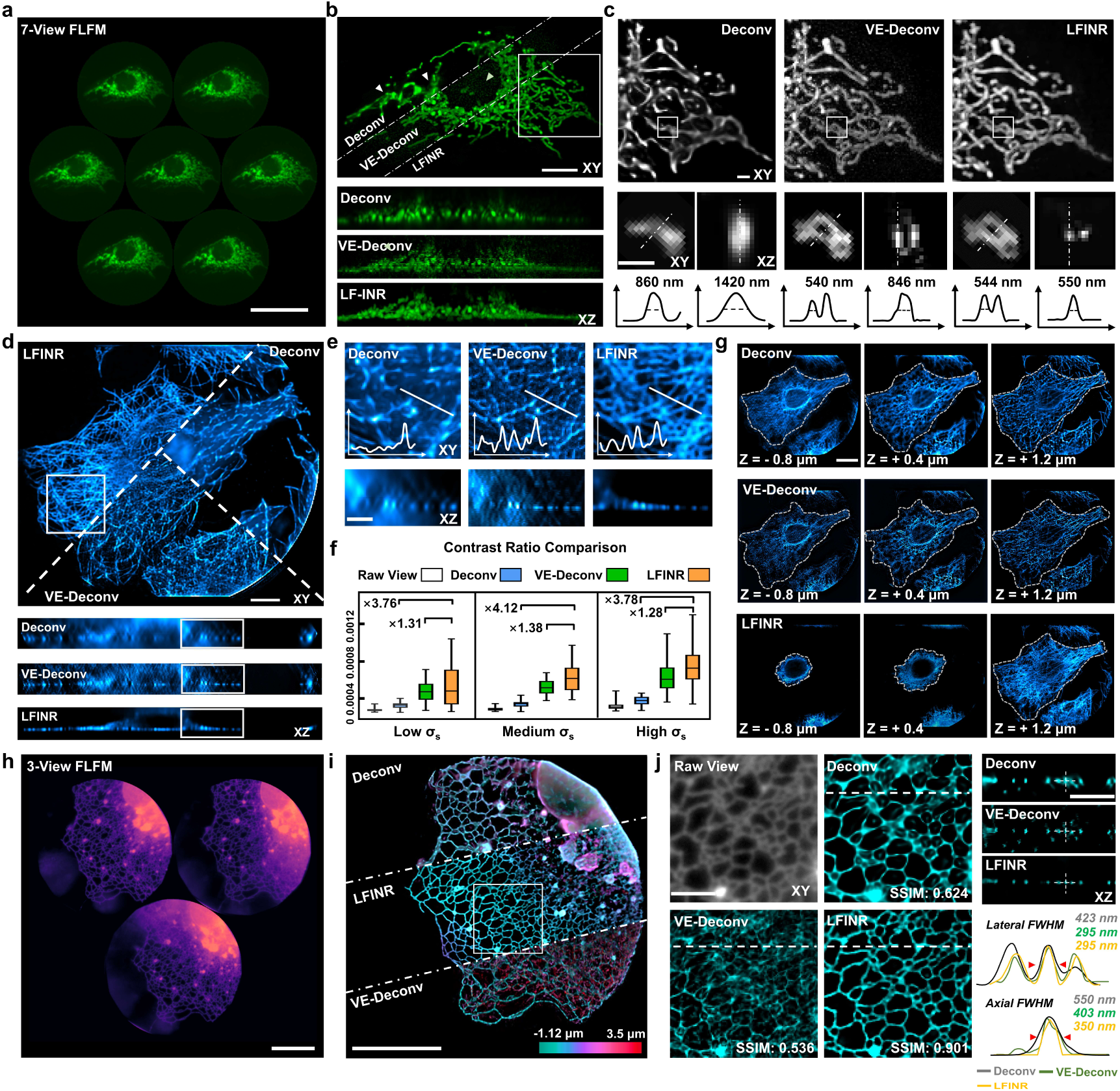
Comparative performance of Deconv, VE-Deconv and LFINR reconstruction on diverse organelles. a, 7 FLFM views of the mitochondria matrix (mito Ma) (tagged with Cox4-EGFP) in a whole live U2OS cell. **b,** The *xy*-MIPs and *xz*-MIPs of mitochondria reconstructed by Deconv, VE-Deconv and LFINR. **c,** The MIPs of Deconv, VE-Deconv and LFINR reconstructions. The magnified lateral and axial views of the ROIs, highlighted by the white boxes, along with intensity profiles along the white dotted lines, show the resolved single mitochondria, highlighting the improved lateral and axial resolutions by LFINR reconstruction. **d,** Comparative results offered by Deconv and LFINR along lateral and axial planes on microtubule (tagged with 3mEmerald- Ensconsin). **e,** The higher-magnification views of microtubule (marked by white rectangular regions in **d**). The plots denote the intensity profiles of the lines shown in the *xy* planes. **f,** The box plots show the statistical contrast ratio^37^ of raw FLFM views, Deconv, VE-Deconv and LFINR results under various signal densities (the deviation of signals: σ_s_). **g**, Comparison of the lateral planes at various axial position reconstructed by Deconv (top row), VE-Deconv (middle row) and LFINR (bottom row). The dotted white line denoted the signal boundary at each depth, showing notably stronger optical sectioning capability by LFINR reconstruction. **h,** The captured 3 FLFM views of ER (tagged with EGFP-Sec61β) in a live U2OS cell**. i,** Depth-encoded *xy* MIPS of 3D reconstructions by Deconv, VE-Deconv and LFINR. **j**, The high-magnification views of white rectangular regions in (**i),** showing notably less hallucinations and axial aliasing. The three insets shown in the right are *x-z* sectional planes along the white dash line in **j**. The intensity profiles along white dotted lines indicated in x-z plane are shown. Scale bars: 50 μm in (**a)**, 10 μm in (**b, d**), 2 μm in (**c**), 5 μm in (**e, j**), 20 μm in (**g, h, i**)

As shown in **Figs. 3a, b,** LFINR reconstruction revealed the detailed structure of mitochondria matrix (mito Ma) while deconvolution results exhibited blurry (denoted by white arrowheads) under 7 views FLFM settings. The line width measurement of resolved mitochondrial in **Fig. 3c** demonstrated that our 7-view LFINR has shown ∼1.58× lateral and ∼2.61× axial resolution enhancement compared to the Deconv results. Moreover, it surpasses the theoretical spatial resolution of FLFM (**Supplementary Table 2**), benefiting from the deblurred views (**Supplementary Note 2**) and the axial enhancement provided by INR (**Supplementary** Fig. 7). This resolution enhancement also improved the structure fidelity of LFINR as shown in **Supplementary** Figure 14. While VE-Deconv effectively reduced the blurring in lateral plans, it still suffered from axial elongation and exhibited more obvious axial artifacts compared to Deconv. Besides, with the robustness in 3D reconstruction with few views, LFINR persevered the structural details while Deconv yielded noticeable artifacts in axial plane (**Supplementary** Figure 15). Furthermore, with the increment of signals’ complexity, the classical model-based deconvolution approach suffered the cross-talk from different slices. As shown in **Figs. 3d**-**e**, the microtube reconstructions by Deconv suffered from the blur and axial elongation, thereby showing lower contrast and couldn’t resolve the hollow structure in cell’s axial slice. VE-Deconv exhibited severe axial artifacts under high signal density. In contrast, LFINR recovered the sharp contrast of microtubules (**Fig. 3e**, **Supplementary** Fig. 14) and enabled 3D visualization of the whole cell. Statistical results (contrast ratio^37^) in **Fig. 3f** demonstrated significant contrast improvements (3.76∼4.12 times higher than Deconv and 1.28∼1.38 times higher than VE-Deconv) across various signal densities. This improved contrast also indicated the superior optical sectioning capability of LFINR in the comparisons of different *xy*- planes in **Fig. 3g**.

In addition, LFINR significantly reduced the artifacts when reconstructing volume from merely three light-field views, under which although the views show higher spatial resolution (**Fig. 3h**), excessive high-frequency hallucinations and cross-talk of depth information also arise (**Fig. 3i**). These undesirable artifacts shown in deconvolution-based results (**Fig. 3i, j**) led to inadequate SSIM metrics and low spatial resolution. In contrast, our LFINR removed the stripe artifacts brought by lower angular sampling rate and achieved enhanced fidelity indicated by higher SSIM values and lateral & axial resolution enhancement (**Fig. 3j, Supplementary** Figure 14).

These high-quality 3D reconstruction with different light-field views verified that LFINR could mitigate the tradeoff between spatial resolution and angular sampling rate and act a powerful inverse-problem solver in multi-view 3D reconstruction.

### Light-field meta neural representation (LFMNR) boosts the 4D visualization of intracellular dynamics

Dynamic biological processes, such intracellular interactions, often occur across three- dimensional space for a long time. High-resolution extraction of the spatiotemporal patterns of these dynamics requires consecutive volumetric reconstructions of light field videos at high throughput, thereby posing big challenge to native LFINR which relies on the scene-wise representation. To make LFINR practical to continuous 4D visualization of biological dynamics, we proposed a weights-transfer strategy based on meta-leaning to enhance the efficiency of sequential 3D reconstruction by inheriting optimized parameters from prior implicit representation network, which was termed as LFMNR (**Fig. 4a, Methods**). Initially, the first frame (LFINR T0) was served as the key-point to perform self-supervised 3D reconstruction with the progressive sampling strategy. Subsequently, the optimized parameters were employed to initialize the network of subsequent LF frame (LFINR T1). Owing to the inherent motion continuity of biological dynamics, the convergence can be reached with merely 100 fine-tuning iterations with hybrid rendering model. As a result, an acceleration of ∼100 folds was achieved as compared to the time consumption on native INR optimization, leading to an ∼20 s quick reconstruction for each volume (**Fig. 4c, Supplementary** Fig. 16). When sequential optimization finished, LFMNR could “memorize” the 4D information in a form of saved network weights. Users could input arbitrary 4D coordinates to query corresponding voxels (**Fig. 4b**) instead of storage of grid-based 4D images. With the unique feature of representation network and meta-learning strategy, our LFINR overcame the deficiency on reconstruction speed, meanwhile it achieved data compression (∼10 folds reduction than the data size of LF views) and computation-cost reduction (∼2 folds) compared with current deconvolution approach (**Fig. 4c**).

**Figure 4.**
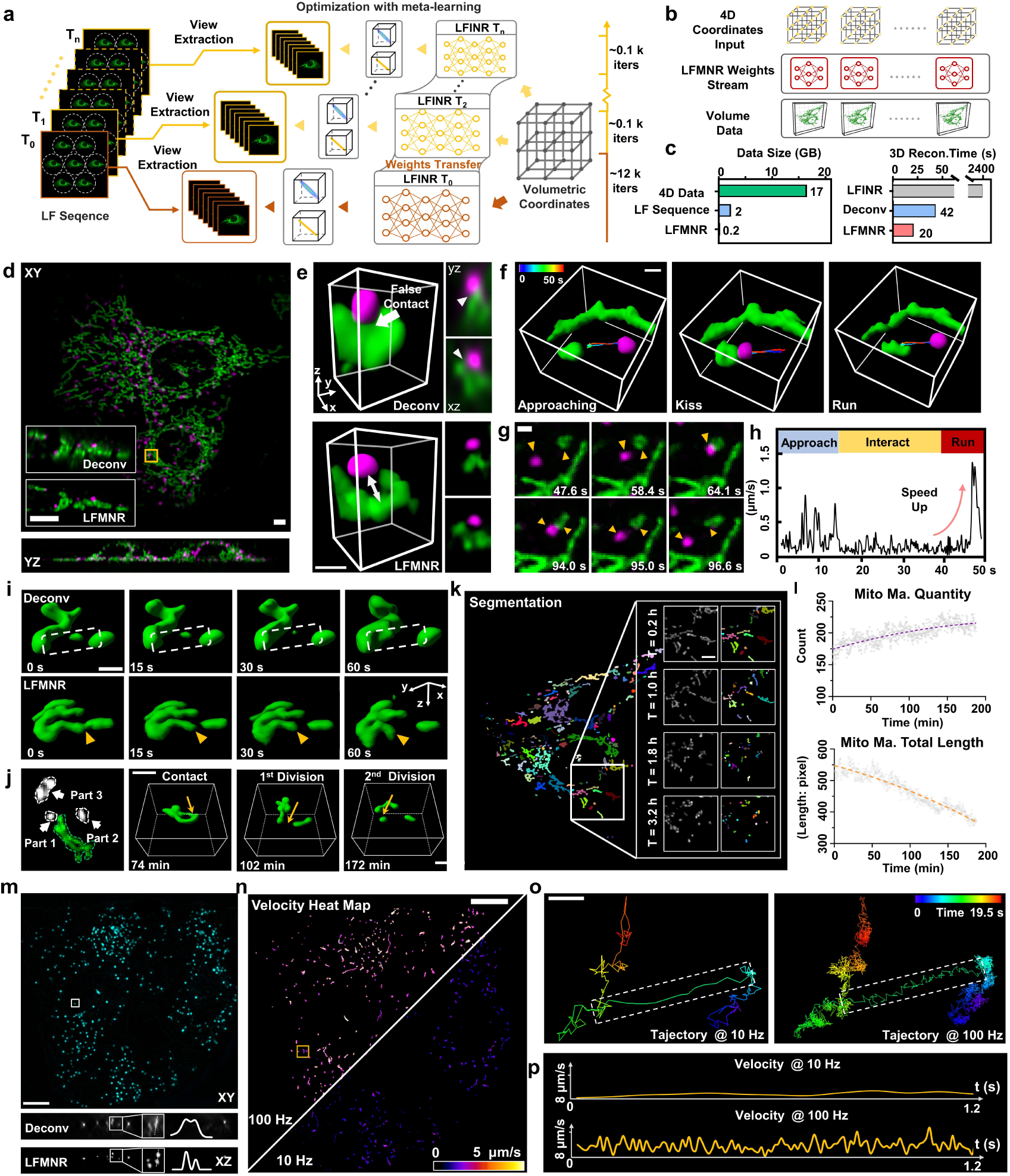
Visualizations and quantifications of intracellular dynamics in live cells by LFMNR. a, The scheme of light-field meta neural representation in solving multi-scenes generalization and accelerating optimization. **b,** The rendering process of 3D volume derived from LFMNR weights. **c,** *Left*: The data size of 4D data (3D videos), LF videos (raw LF sequences) and LFMNR weights (network parameters), denoting the strong data-compression ability of LFMNR. *Right*: The time- consumption comparison of Deconv and LFMNR for 3D reconstruction of light-field videos. **d,** Dual-color imaging of mitochondria (Green, tagged with Cox 4-EGFP) and lysosome (magenta, tagged with Rab7-mCherry2) in live U2OS cells. Insets show the comparison of axial section between Deconv and LFMNR. **e,** The volume renderings and slices of the regions of interest (ROI) indicated by orange boxes in **d**, demonstrating more accurate identification of mitochondria- lysosome contacts enabled by higher resolution of LFMNR compared to Deconv. **f,** The volume rendering of the interaction process between mitochondria and lysosome occurring in 50 seconds. **g,** The lateral projection of dual-color LFMNR results illustrating the fast “kiss” (top panel) and “run” (bottom panel) process. **h,** The velocity changes of lysosome during the whole process visualized in **f**, quantifying the phenomenon of accelerated running away of lysosomes after detaching with mitochondria. **i,** LFMNR reconstruction of the first frame of the long-term mitochondria evolution during cell apoptosis. The ROI show more accurate identification of mitochondria fission events achieved by higher fidelity of LFMNR compared to Deconv. **j,** The dynamic process of mitochondria over an observation period of 3.3 hours. **k,** The segmentation result of LFMNR reconstruction show the fragmentation of mitochondria during observation. **l,** The statistical results of the total length and quantity of mitochondria during the whole observation based on the segmentation approach in (**k)**. **m**, x-y and x-z MIPs of peroxisomes (tagged with SKL- mApple) in a live U2OS cell, imaged at a speed of 100 Hz and reconstrued using LFMNR and Deconv. **n,** Velocity heat map tracking the 3D motion of peroxisomes at 10 Hz (left) and 100 Hz (right). **o,** Motion trajectory of a selected single peroxisome (orange box in **n**) captured at 100 Hz and 10 Hz, over a 19.5 s duration. **p,** The corresponding velocity plots of the trajectory highlighted by white dashed box in **o**. Scale bars: 5 μm in (**d)** and the insets in (**k**), 1 μm in (**e, f**), 2 μm in (**g, i, j**), 20 μm in (**k**), 10 μm in (**m, n, o)**.

To demonstrate the utility of LFMNR, we conducted dual-color live-cell imaging to capture the locomotion and interaction of mito Ma (tagged with Cox4-EGFP) and lysosome (Rab7-mCherry2) in three dimensions, as shown in **Fig. 4d**. We visualized the instantaneous 3D interactions between two types of organelles (**Figs. 4e**-**g**) at 10 Hz volume rate. LFMNR enabled precise delineation of the distance between mitochondria and lysosomes, whereas Deconv results incorrectly indicated a mitochondria-lysosome contact due to axial elongation and reconstruction artifacts (**Fig. 4e**). With LFMNR’s accurate visualization, we tracked lysosome velocity during the “approaching- contacting-separating” process and identified a dramatic speed increase as lysosomes moved away (**Figs. 4f**-**h****, Supplementary Video 1**). Besides the fast interaction between organelles, we also conducted long-term observation (∼3.3 hours) of mitochondria evolution (**Figs. 4i, j, Supplementary Video 2**). LFINR successfully captured multiple mitochondria division events and tracked the resulting daughter mitochondria completely, whereas Deconv exhibited signal loss, failing to detect mitochondrial fission events (**Figs. 4i**). In addition, leveraging the high structural contrast of LFINR results, we segmented and identified each mitochondrion, revealing the smaller and rounder mitochondria resulted from extensive fragmentation of mitochondria shape (**Fig. 4k, Supplementary Video 3**). The statistical variation of mitochondria length and quantity in **Fig. 4l** also demonstrated the mitochondrial fragmentation process, indicating the potential apoptosis phase of the cells^38, 39^. The long-term observation capability enabled by the highly efficient photon utilization and high-fidelity reconstruction of LFMNR allowed for the visualization of cell mitosis over 25 hours (**Supplementary** Fig. 17).

Furthermore, we imaging the fast motion of peroxisomes in live U2OS cells at a volume rate of 100 Hz (**Fig. 4m, Supplementary Video 4**). While Deconv failed to resolve peroxisomes due to low axial resolution and artifacts, LFMNR successfully resolved these subcellular structures in three dimensions, enabling the extraction of high spatiotemporal resolution patterns of peroxisomes motion (**Fig. 4n**). In contrast, tracking the dynamics at 10 Hz, the maximum volumetric imaging rate achievable by current scanning-based microscopes, resulted in significantly less accurate trajectory and velocity quantifications (**Fig. 4n-p**).

### LFMNR enables 4D reconstruction and quantification of instantaneous functional signaling activities in live zebrafish larva and *C.elegans*

LFMNR is also capable of recording and analyzing *in vivo* dynamics occurring in three dimensions within the live tissues.

We first capture the calcium dynamics (labeled with *huc: gcamp6s*) across the whole brain of an awake zebrafish larva at 10-Hz volume rate^40^ (**Supplementary Video 5**). While the Deconv results showed ambiguous neuronal structures due to its limited resolution, LFMNR effectively resolved the complex structure of the whole brain at near single-cell resolution over a large tissue depth of 240 µm (**Fig. 5a, b)**. As a result, LFMNR provided a clearer identification of calcium signal fluctuations compared to Deconv (**Fig. 5c**).

**Figure 5.**
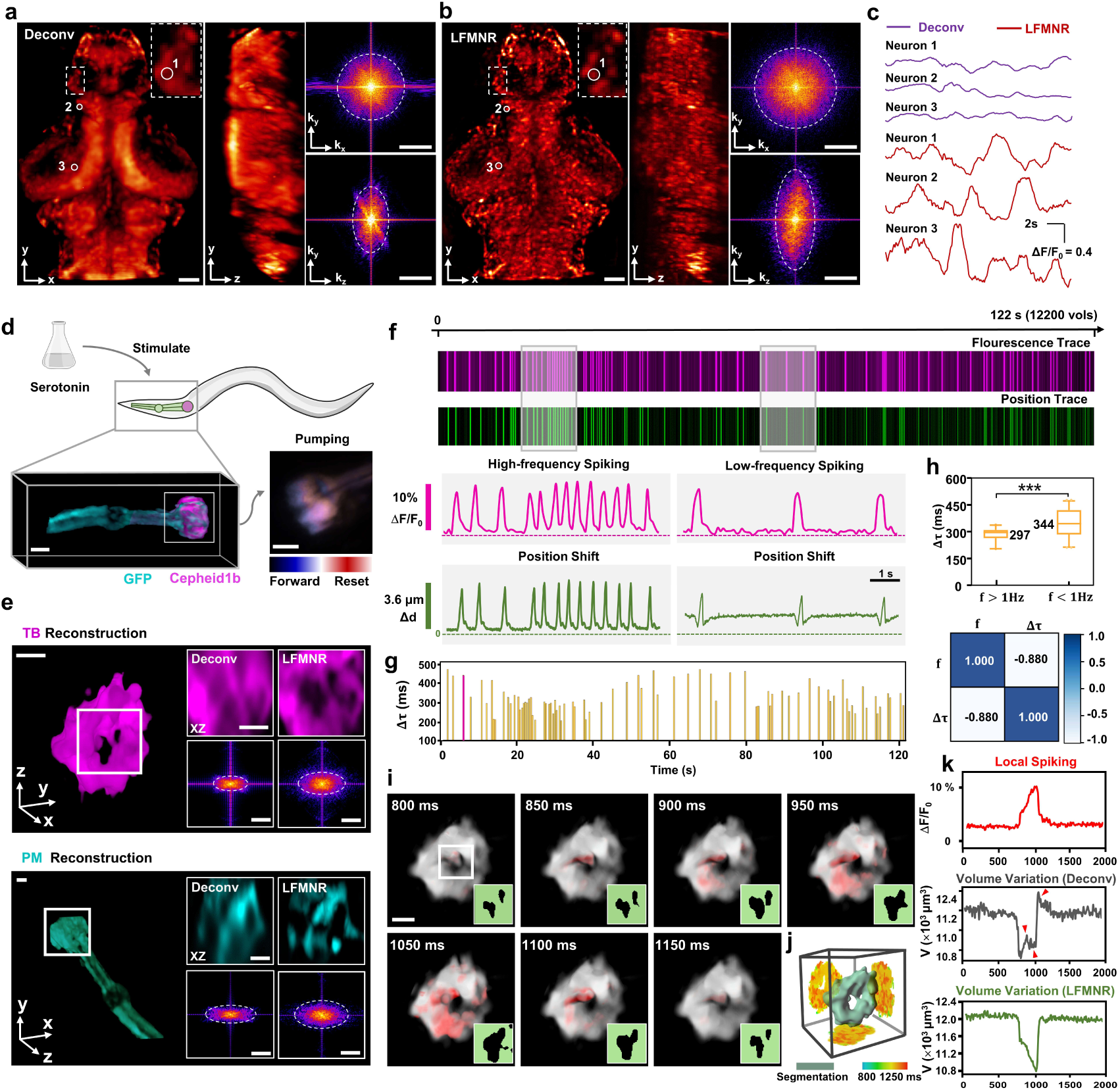
LFMNR-based visualization and quantification of instantaneous functional signals in behaving zebrafish and *C*. *elegans*. a, b, Orthogonal MIPs of zebrafish brain with pan-neuronal cytoplasm-labeled GCaMP6s (huc:gcamp6s) reconstructed by Deconv and LFMNR. The insets show the magnified view of the ROI indicated by the white boxes. **c,** Calcium dynamics of the same neurons obtained by Deconv and LFMNR. **d,** Dual-color light-field imaging of pharyngeal muscles in C. elegans fluorescently-labeled with Cepheid1b and GFP. The temporal-encoded LF captures (lower right) indicate the pharyngeal pumping dynamics stimulated by the serotonin (**Methods**). **e**, The comparison between deconvolution results and LFMNR reconstructions of the pharynx. TB: terminal bulb. PM: pharyngeal muscle. **f**, The variation curve of voltage signal intensity and centroid position extracted from the LFMNR reconstructions of Cepheid1b and GFP channels, respectively. Over 12000 consecutive volumes were reconstructed during 122-second observation. The spikings shown in the two curves denoted the fluorescence fluctuation and position shift of *C. elegans*, respectively, during the pumping process. The time windows marked by the boxes show two typical spiking traces with different frequencies. **g**, The plots of spiking width (hτ) extracted from whole fluorescence trace in **(f)**. **h,** Top: Bar chart of hτ under two spiking frequencies. n>10. *P*= 0.0005, Mann-Whitney test. Bottom: The heatmap of correlation matrix calculated from spiking frequency and spike width. *f*: frequency. **i**, The reconstructions of terminal bulb (*F*, gray scale) and its voltage signals (*F-F_0_*, red) during a single spiking event highlighted with magenta peak in **(g)**. **j**, The segmentation of terminal bulb (green) and temporal-encoded projections revealing voltage signal propagation in 3D space during a 450-ms spiking event shown in **(i). k**, The tightly-synchronized peaks between pharyngeal lumen dynamics and voltage signal spiking. Scale bars: 50 μm in (**a, b**), 15 μm in (**d**), 10 μm in (**e, i**). Scale bars in Fourier spectrum images: 1/8 μm^-1^ in (**a, b**), 1 μm^-1^ in (**e**)

In addition to the volumetric speed requirements of functional calcium imaging, understanding how membrane voltage mediate behavior requires ultra-high-speed recording^41, 42^. The transient voltage signals within the functional activity occurred irregularly at different 3D positions and accompanied with local morphology changes. While LFM has become an optimal choice for capturing the 4D dynamics of functional voltage signals, the high-resolution reconstruction of such instantaneous signals with unknown structures poses significant challenge to either classical model based^9, 10^ or volume-supervised LFM approaches^15, 17, 19^. In contrast, LFMNR demonstrated timelapse imaging of the pharynx of *C*. *elegans* (labeled with Cepheid1b^43^ and GFP) to capture the electrophysiological activities and corresponding 4D voltage signals distribution during the pumping process of worm, with 100-Hz volume rate (**Fig. 5d**). While conventional deconvolution results suffered from severe axial blur, LFMNR reconstructed pharynx with higher structural contrast and better resolution to distinguish the pharyngeal lumen (**Fig. 5e**). Meanwhile, the efficient meta-learning enabled sustained reconstruction (12200 volumes) of the transient voltage spiking and 3D locomotion of labelled signals during pharyngeal pumping (**Fig. 5f**). Each peak in the position trace curve (magenta line) represented one ‘forward-reset process’ by the contraction and relaxation of pharynx muscle, consistent with the voltage-induced fluorescence fluctuation (magenta line). From the reconstructed fluorescence traces, we identified high-frequency and low-frequency voltage spiking corresponding to diverse modes of locomotion. We further extracted all the voltage peaks across entire 120- second observation time and quantitatively analyzed the width (hT) of each peak (**Fig. 5g, Methods**). We found that the peak width exhibited a strong negative correlation with the spiking frequency, showing a significant disparity in hT between high- frequency spiking >1 Hz and low-frequency spiking <1 Hz (**Fig. 5h**). In addition to the mapping of various spiking events, we specifically visualized the voltage signaling during one event. The sequential LFMNR reconstructions (**Fig. 5i and Supplementary Video 6**) detailed the morphological changes of pharyngeal lumen (gray scale with white rectangular), which were accompanied with local transient voltage spikings (color-coded with red). After continuously segmenting the LFMNR reconstructions, we further illustrated how the transient 3D voltage signals propagate in 3D space over a temporal scale of 450 ms (**Fig. 5j**). The enhanced spatial resolution achieved by LFMNR enabled precise synchronization of the spiking process with the volume changes of the pharyngeal lumen. In contrast, the volume changes segmented from Deconv reconstructions exhibited irregular fluctuations (red arrows) due to artifacts and low axial resolution. (**Fig. 5k**).

## Conclusion and Discussion

Achieving high-structural-fidelity 3D reconstruction in light-field microscopy, especially when the angular captures are limited and sparse, is essential for accurately capturing dynamic biological processes at high spatiotemporal resolution. Classical model-based approaches are inefficient due to the absence of prior knowledge. While supervised-learning methods offer a solution by constructing a “2D-to-3D” mapping function guided by volumetric data, they are laborious and costly in preparing labeled datasets and often suffer from poor generalization. Furthermore, the recovery of dense signals remains a challenge to all current light-field reconstruction approaches.

Through the combination of continuity bias of INR model and physics-based hybrid-rendering model, our LFINR approach can achieve high-throughput and high-quality 3D reconstruction from a single 2D FLFM capture in a completely self-supervised manner. Compared to conventional deconvolution approach, LFINR eliminates axial artifacts and elongation for dense signal 3D reconstruction, maintaining high structural fidelity with low errors. Notably, even in scenarios involving few-views reconstruction, LFINR maintains high structural fidelity due to its inherent geometric consistency across predicted views. INR’s view-synthesis ability also mitigates axial elongation caused by the missing cone problem, enabling artifact-free 3D reconstruction and achieving spatial resolution enhancement (>1.4x) across various samples, from subcellular structures to *in vivo* tissues. Furthermore, we extend the LFINR framework to OPT, FPM and Cryo-ET techniques, demonstrating its high versality in addressing inverse tomographic reconstruction problems (**Supplementary** Figs. 10-13).

In addition to the improvement of reconstruction quality, to mitigate the computational cost in conventional uniform-random sampling strategy, we propose the progressive sampling with the aid of structural error map to accelerate convergence speed and prevent the signal loss. Furthermore, to make LFINR practical in visualizing sustained biological dynamics, we develop a meta-learning approach to expedite network convergence through temporal correlations, achieving ∼two-order-of-magnitude acceleration with averaged inference time of merely 20 seconds for one volume. Through the self-supervised transfer-learning strategy, LFMNR can visualize the 5D (3D space + time + spectrum) intracellular events with higher structure fidelity compared with classical model-based approach, and record the transient 4D (3D space + time) voltage signals that are challenging to both classical model-based or volume- supervised LFM approaches.

In summary, our LFMNR utilizes neural networks to smoothly represent high-dimensional scenes (3D volumes) with low-dimensional observations (2D FLFM views). This new “scene representation” paradigm solves the issue of inadequate quality in classical model-based light-field reconstruction approaches while completely eliminates the dependence on data priors. Meanwhile, it not only enhances the image quality for light-field 3D reconstruction but also exhibits capability in data compression. We envision that LFMNR can be further optimized through the future efforts on training acceleration for real-time reconstruction^44, 45^, combination with high-quality optical setups for higher resolution beyond diffraction limit of the views^46^, and robust restoration under diverse degradation models^47, 48^.

## Data availability

We provided representative numerical simulation data and experimental data, which are available at https://github.com/feilab-hust/LFMNR. All other data that support the findings of this study are available from the corresponding author upon request.

## Code availability

All the codes of LFMNR and corresponding usages can be found at https://github.com/feilab-hust/LFMNR.

## Acknowledgments

This work was supported by the funding from the National Natural Science Foundation of China (T2225014, 62375095, 21927802), National Key Research and Development Program of China (2022YFC3401100, 2023ZD0519900). The authors are grateful to Longbiao Chen and Yao Zhou for their helpful discussion, and Guodong Tan and Quan Wen for their provided experimental data.

## Author contributions

P.F. and C.Y. conceived the idea. P.F., B.L., D.L. oversaw the project. P.Z., G.T., Y.L., S.G., Q.W., M.Z., and Y.Z. provided the samples. J.S., C.Y. and L.Z. developed the optical setups and acquired the experimental images. C.Y., S.W., M.S. and Y.M. developed the programs and processed the images. Y.L. and M.Z. provided the samples and conducted the biological experiments. C.Y., M.S., L.Z., Z.W., S.G., T.K.H., Y.Z., B.L., L.Z., D.L. and P.F. discussed and wrote the paper.

## Methods

### FLFM optical setup

The FLFM system was built on a commercial inverted microscope (IX73, Olympus). To demonstrate the versatility of LFINR under various hardware settings, we conducted imaging experiments with both 3-view, 7-view and 19-view configurations.

For the 3-view setting in cell imaging, a 100x/1.4 objective lens (Olympus UPLSAPO100XO/1.4) was used to collect fluorescent emission, which was controlled by an objective scanner. To prevent the overlap between captured views, an iris (SM1D12, Thorlabs) was positioned as the native image plane. The focal length of Fourier lens (FL) was set to 250 mm. A microlens array (*pitch* = 3.25 mm, *f-number* = 37, *fML* = 120 mm) was placed at the back foci plane of Fourier lens, producing 3 FLFM views. Then, the multi-view signals were collected by an sCMOS (Prime BSI Express, Teledyne Photometrics), producing the effective FOV (field of view) of 67.7 μm. The whole schematic of the optical path is shown in **Fig. 1a**.

For the 7-view setting in cell imaging, we adopted a FLFM configuration with a 60x/1.5 oil immersion objective lens (Olympus UPLXAPO60XO) and a Fourier lens (*fFL* = 250 mm) to achieve Fourier transformation and cover 7 microlens. The microlens array and camera were the same with 3-view setting. The FOV was 112.85μm.

For the 7-view setting in pharynx muscle imaging, a 30x/1.05 objective lens (Olympus UPLSAPO30XSIR) and a Fourier lens with 250 mm focal length were used. The pitch and focal length of microlens array were 3.63 mm and 120 mm. The FOV was 252.08 μm.

For the 19-view setting in voltage imaging on terminal bulb, a 30x/1.05 objective lens (Olympus UPLSAPO30XSIR) was used to collect fluorescent emission. The focal length of Fourier lens is 250 mm. The pitch and focal length of microlens array is 3.25 mm and 120 mm, respectively. To capture the fast voltage signals, we used a high-speed sCMOS (Kinetix, Teledyne Photometrics) and obtained 19 views with a FOV of 225.69 μm,

The optical parameters of numerical simulation in **Fig. 2** are also included 3-view 7-view and 19 views settings. The corresponding system parameters are listed in **Supplementary Table 2.**

### 3D reference data generation

To quantify the performance of different reconstruction method on various biological samples in, we used scanning-based microscope to acquire high-resolution 3D data as the reference 3D data in simulation experiments in **Fig 2**. Specifically, in **Fig 2b**, the microtubule data was acquired via a self-built Oblique Plane Microscopy^49^ (OPM) using an Olympus ix83 microscope. The primary objective was a 60×/ 1.3 silicone oil objective (UPLSAPO60XS2, Olympus). In **Fig. 2d**, the 3D ER (endoplasmic reticulum) data came from the confocal microscope (ZEISS LSM 980 with Airyscan2). To obtain the 3D cardiomyocyte nuclei data in **Fig. 3g**, we conducted confocal imaging (SP8- STED/FLIM/FCS, Leica) using a 20×/0.75 W objective (HC PL APO CS2) on deeply anesthetized fish larvae. After obtaining the raw 3D stacks, we first conducted background subtraction in Fiji to remove the uneven background. Then, these stacks were resampled following the sampling rate of various FLFM optical setups. Such resampled stacks were used to generate FLFM projections with simulated PSF, and also served as the “GT” images to assess the reconstruction quality of various approaches.

### Progressive sampling strategy

INR-based approach^24^ adopted random sampling strategy to constant optimize the scene-representation network. The number of coordinates, which increase exponentially as the 3D scene size increases, affects the network convergence speed. Further, when applied such random sampling strategy to represent the 3D biological scene, the uneven distribution of grayscale intensity and structural complexity in fluorescent images made this representation difficult, denoted by the weak-signal loss and slow convergence (**Supplementary** Fig. 3). We attributed this deficiency to the ignoration of each voxel’s contribution when conducting loss summation and averaging, implying that the network tended to repeatedly optimize the “easy” area with low-frequency and high intensity^50^. Inspired by the hard example mining in object detection^51^, we developed a progressive sampling strategy to focus on the positions where contains structural distortion and signal loss.

The sampling strategy contains 3 steps: sparse sampling, full-sampling and focal sampling as shown in **Fig. 1a**.

Firstly, for the optimization process of one scene, the captured FLFM views were down-sampled to half the size of original ones. Accordingly, the spatial meshgrid and the projection matrixes (FLFM PSF and geometry projection matrix) in hybrid physics model were also re-sampled to match the location and sampling rate of the resized views. Based on the low-frequency implicit bias^50^ in neural network, this sparse sampling strategy simplified the fitting task on complex 3D signal and reduced the number of query voxels, thereby promoting the convergence at the initial stage.

Secondly, the neural network was optimized under the supervision of multi-views with the original sampling rate to recover the high-frequency details of the 3D scene, which would exist signal loss like random sampling strategy. Therefore, during the optimization process, an error map was simultaneously calculated and saved for the following step as the sampling probability map. Denoting *^N^*^×*N*^ as the center- view projection from LFINR 3D prediction with geometry-model, and *^N^*^×*N*^ as the target enhanced center view. The error map can be formulated as:

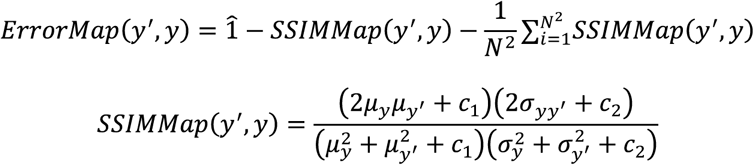

where means an all-ones matrix with the size of center view and *N* means the lateral size of the center view. *_y_* and *_y_′* are the mean of and, respectively. *^2^* and *^2^* stand for the variance of and *y*. *y^2^* and *^2^* are constants for 32-bit data (normalized to (0,1)). The kernel size used in SSIM calculation was set to 7. Then, we converted this map into a sampling probability map via dividing each pixel value by the summation of error map.

Thirdly, based on the calculated dynamic probability map, we conducted selective optimization on different regions of scene to recover the structural details. The time consumptions of various samples were listed in **Supplementary Table 1**.

### Hybrid model rendering

During the view rendering phase, LFINR adopted hybrid physics model to generate dual forward projection. The first term in the hybrid model was described by the FLFM PSF based on the wave-optics theory. In this paper, we modified the source code of oLaF^52^ to generate PSF matrix, which was then convolved with represented 3D scene to produce wave-optics projection. The second term involved a geometric-optical model to describe the propagation process of ideal rays. Similar to the convolution process in first part, this rendering process can be also described as the convolution between 3D volume and geometry projection matrix. To obtain this matrix, the 3D positions of maximum value of FLFM PSF at different depth were extracted first. Then, conducting linear fitting on these coordinates to obtain the depth-wise shift functions of different views:

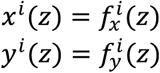

where *i* is the view index and the 3D spatial coordinates. *^i^* and *_i_* are the fitted parallax function, which revealed the projective geometry relation. Then, input all -coordinates into these two functions to obtain the 3D positions of chief rays. Finally, an all-zero matrix with the size of the FLFM PSF was created, and voxels at the obtained 3D positions were assigned a value of 1.

### Geometry consistency

To assess the fidelity of 3D reconstructions and underscore the view-synthesis ability of LFINR in Fig. 2, we employed geometry-consistency, the structural similarity of Epi-polar Plane Images (EPIs)^30^ computed from 3D volume. The geometry-consistency can be formulated as:

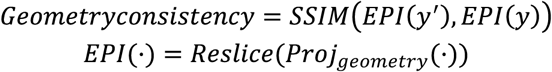

where *y’* is the reconstruction result from deconvolution-based methods or LFINR and is the 3D ground truth. means computing the EPIs from 3D volume. *_geometry_* refers to the generation of views with horizontal arrangement by a tomography projection model. The number of generated views was set to 19 to ensure angular continuity. The angle range of this model was set to *_objobj_* where *_obj_* is the maximum acquisition angle decided by objective’s NA. The operation of *Reslice* involves two steps: *a*. Concatenate the generated 2D projections along z-axis to obtain “view-stack”; *b*. “Top-reslice” the “view-stack” in *Fiji* to obtain axial section. This section plane is the EPI, containing both spatial information and angular information.

The whole process is illustrated in **Supplementary** Fig. 9. **Network implementation of LFINR**

During one network optimization process, the input coordinates would follow 3 steps to obtain the computed loss: 1) Position encoding 2) MLP inference 3) Loss computation (**Supplementary** Fig. 2).

***Position encoding***. The coordinates’ low-frequency variation among 3D space would degrade the rendering quality of structural details or color variation^24^. Therefore, in LFINR, we adopt the Fourier positional encoding to convert such variation to the encoded high-frequency information:

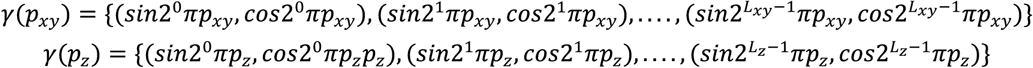

where *_Xy_* is the lateral coordinate (*x*, *y*) while *_z_* is the axial coordinate (*z*). *_Xy_* and *_z_* are lateral and axial encoding order, respectively. The detailed settings of *_Xy_* and *_z_* in various sample are listed in **Supplementary Table 1**. After position encoding, a high-dimensional feature vector would be obtained, denoting as *^N^*^×*L*^, where *N* is the number of input coordinates and *L* is the channel of encoded feature deiced by *_Xy_* and *_z_*. Each element *_i_* in can be expressed as:

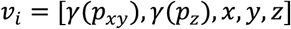

***MLP inference*.** We utilized a Multi-layer Perceptron (MLP) to represent the 3D scene with querying spatial coordinates. MLP consists of input layer, hidden layers and output layers. The detailed parameters are seen in **Supplementary Table 1**. This process can be formulated as:

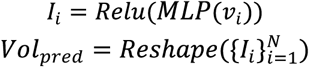

where is the index of some voxel and is corresponding grayscale. is used to truncate the predicted values smaller than zero. After network inference, the collection of predicted voxels would be reshaped to the 3D-grid image (*_pred_*).

***Loss computation***. In LFINR, the loss function consisted two parts: L2 loss function in wave-optics projection process and structural loss function in geometry projection process (**Fig. 1a**). Specifically, the represented volume was convolved with FLFM PSF to fit the captured LF views and also with geometry projection matrix to fit enhanced views. The objective function of network optimization based on these views can be expressed as:

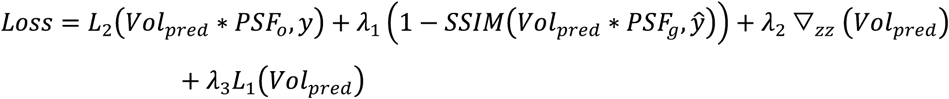

where *_o_* and *_g_* are wave-optics PSF and geometry PSF, respectively. and are enhanced views and raw views. *_zz_* and *_1_* are the regularization term to avoid model overfitting. *y_1_*, *_2_*, *_3_* are weighting coefficients (**Supplementary Table 1**).

### Data preprocessing

In this study, experimental LF captures were pre-processed to remove the extra image degradation derived from fluorescent background or noise. For static scene, the raw LF images underwent background subtraction by rolling-ball subtraction in Fiji. Subsequently, the mean intensity of each view was rescaled to be the same. For dynamic scenes, we used rolling-ball background subtraction on LF captures of live cells and subtracted a constant background from the voltage LFs. Due to the noise inherent in *liv*e cell imaging and voltage imaging, a lightweight self-supervised denoising network (**Supplementary Note 4**) was constructed to enhance the signal-to-noise ratio (SNR) based on the captured LF videos.

### Classical model-based algorithm

To evaluate the performance of self-supervised LFINR, we selected the classical model-based algorithm, Fourier light-field microscopy deconvolution^52, 53^ (Deconv), as the baseline approach. The FLFM point-spread function (PSF) used and the 3D reconstruction algorithm in this study derived from the open-source code^52, 54^, based on the wave-optics theory. For different FLFM setups, PSF simulations were conducted according to the optical parameters listed in **Supplementary Table 2**. The number of iterations was 30.

Besides, to further highlight the advantages of LFINR, we developed the “view- enhanced deconvolution” (VE-Deconv) method. In VE-Deconv, the enhanced FLFM views, produced by the self-supervised view-enhancement network (**Supplementary Note 2**), were used to yield 3D reconstructions instead of the raw FLFM views. Accordingly, the PSF was replaced with the geometric projection matrix for each optical setup, while the number of iterations was also 30.

### Voltage signal analysis

In voltage signal analysis shown in **Fig. 5**, we calculated the position shift and fluorescence trace from the 4D reconstructions of terminal bulb (TB). For the calculation of position shift, we first extracted the binary masks of 3D volumes and recorded the 3D position of masks’ centroids. Then, calculate the distance of extracted positions relative to the position of *C*. *elegans* under resting-state. The fluorescence traces were calculated by Δ*F*/*F*0 = (*F* − *F*0) / *F*0, where *F*0 is the mean intensity in the reconstructed TB signal averaged over the entire time series, and *F* is the sum intensity of the TB signal. Then, the trace was analyzed for peak extraction by calculating the local maximum within a sliding window of 300 ms, step size of 100 ms. The location of local maxima that surpassed the predefined threshold was recorded as the timepoint of one spiking event. The threshold was set as 10% of the maximum value of the fluorescence trace. Furthermore, the trace was analyzed to extract start and end of peak by searching for the first and last data point to cross a line at 10% of the distance from the baseline (*F*0) to the peak maxima, respectively^55^. The distance between searched points was defined as the spiking width.

### Dynamic-scene representation by LFMNR

The essence of implicit neural representation (INR) is to represent complex high-dimensional signal smoothly with neuron network. For living biological imaging, the signal varies along 3D spatial coordinates and a 1D time axis, presenting challenges for network representation. One plausible approach is to add the ‘*t*-variable’ directly into INR optimization to learn the 4D function . However, this approach significantly increases the computational burden due to the massive inputs of four-dimensional coordinates, especially for long-term imaging. Moreover, sudden grayscale changes caused by fast motion further complicate the four-dimensional function. Another approach involves employing an additional network to predict the motion modality of 3D signal^56^, but constructing such motion model requires sophisticated handcrafting, making it challenging to accurately model irregular motion patterns and local morphologies of biological samples. Inspired by redundancy elimination techniques in data compression^57^, we proposed a meta-learning-based approach to represent dynamic 3D scenes encoded in light field sequence, termed light-field meta neural representation (LFMNR), enabling the acceleration of network convergence based on high inter-frame correlation. Meta-learning, often referred to as “learning to learn”, is a broad concept in metacognition that involves understanding and improving one’s own learning processes. In this context, meta-learning refers to using optimization results from one scenario as the starting point for the next scenario’s optimization, aiming to accelerate the learning process of neural networks. Specifically, during training phase, we first optimized network to represent the first volume with the guidance of input T1 views, which produced network weights *y* and optimizer *y_1_*. The learning rate was 5×10^-4^ and the number of total iterations were 12000. For the second volume, the network and optimizer were initialized with parameters *y* and with *_1_*, respectively. Then, the network was updated under the supervision of T2 views. For remained volumes, according to consecutively transferring the previous parameters, the network convergence time was dramatically. During transfer-learning stage, the number of iterations was 100 for each time-point, which cost ∼20 s.

1. *C. elegans* strains maintenance and generation

All strains of *C. elegans* were cultured on standard Nematode Growth Medium (NGM) plates, which were seeded with OP50 and consistently maintained at a temperature of 22°C^58^. Unless otherwise stated, the Bristol N2 strain is the wild-type animal in reference. Hermaphrodite adults, specifically those that were one day old, were utilized in all the conducted experiments. Transgenic worms were produced employing standard microinjection techniques, wherein 40 ng/μl of specific plasmid DNA was injected^59^.

### Molecular biology

The manipulation of Cepheid1b’s expression^43^ within the *C. elegans* was achieved through the employment of the Three-Fragment Multisite® gateway system (Invitrogen™, Thermo Fisher Scientific, Waltham, MA, USA)^60^. More specifically, three entry clones were utilized: Slot1, Slot2, and Slot3, which were corresponded respectively to the promoter, the target gene, and the fluorescent marker gene. They were subsequently recombined into the pDEST™ R4-R3 Vector II, initiating a LR reaction that resulted in the formation of expression plasmids^61^. For the specific purpose of driving the expression of Cepheid1b within the pharyngeal muscles of *C. elegans*, the *myo-2* promoter was used. This experiment also incorporated green fluorescent protein (GFP) that functioned as a control marker.

Voltage imaging in *C. elegans* pharyngeal muscle.

The day prior to the voltage imaging experiment, *C. elegans* in the L4 stage were relocated to standard NGM plates (35mm in diameter, Nantong Baiyao Laboratory Equipment Co., Ltd.) which were seeded with an OP50 bacterial suspension (OD600=0.9) . This suspension was accompanied by the addition of ATR (1 mM, Sigma-Aldrich, USA) to enhance the brightness of Cepheid 1b. To stimulate pharyngeal pumping, a volume of 20 µl of serotonin was introduced into the recording bath solution (final concentration to be 30mM) for an interval of 3 minutes prior to the commencement of the experiment. The worms were then gently affixed (using Histoacryl Blue, Braun, Germany) to a 2% agarose-coated cover glass, which was immersed in the bath solution. The solution was composed of the following elements (in mM): NaCl 150; KCl 5; CaCl2 5; MgCl2 1; glucose 10; sucrose 5; HEPES 15, pH 7.3 (adjusted with NaOH), ∼330 mOsm.

### Cell culture

U2OS cells were grown in culture medium containing McCoy’s 5Amedium (Thermo Fisher Scientific) supplemented with 1% antibiotic-antimycotic (Thermo Fisher Scientific) and 10% fetal bovine serum (Thermo Fisher Scientific) at 37 ℃ with 5% CO2 in a humidified incubator.

For labeling mitochondrial matrix and tubulin in live U2OS cells, cells were first transfected with Cox4-EGFP and 3mEmerald-Ensconsin using Lipofectamine 2000, respectively, according to the standard protocol and cultured at 37 °C with 5% CO2 for an additional 8 hours. After 6-8 hours of transfection the cells were digested with 0.25% trypsin, seeded on cell culture dishes (20 mm diameter), and incubated for 36 hours at 37°C in 5% CO2.

For labeling lysosomes in live U2OS cells, cells were first transfected with Rab7A- mCherry2 using Lipofectamine·2000 according to the standard protocol and cultured at 37°C with 5% CO2 for an additional 24 h. Before imaging, remove old media and add fresh media.

For labeling ER in fixed cells, U2OS cells were first transfected with Sec6lβ-EGFP for ER using Lipofectamine 2000 according to the standard protocol and cultured at 37 °C with·5%·CO2 for an additional 24 hours. For fixed cells imaging, the cells were fixed with 2% glutaraldehyde for 20 mins.

For labeling peroxisomes in live U2OS cells, cells were first transfected with SKL- mApple using Lipofectamine·2000 according to the standard protocol and cultured at 37°C with 5% CO2 for an additional 24 h. Before imaging, remove old media and add fresh media.

