## Supplementary Materials for "High-fidelity generalizable light-field reconstruction of biological dynamics with physics-informed meta neural representation"

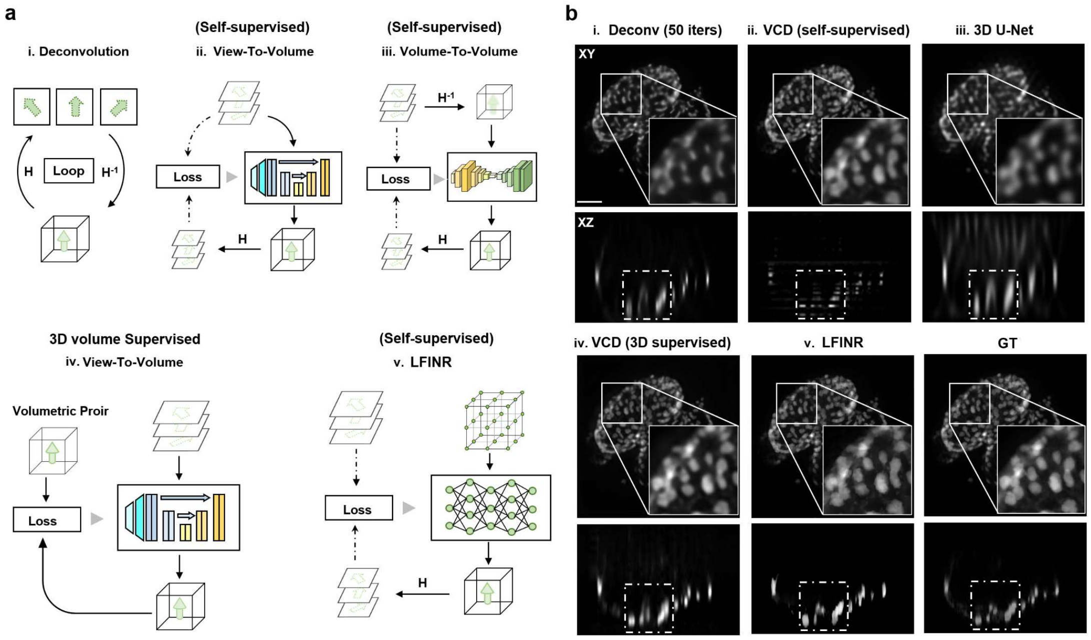


**Supplementary Figure 1. Comparison of various model based and deep-learning based paradigms for 3D light-field reconstruction**. **a,** Principle of different 3D reconstruction paradigms. i, Deconv: Richard-Lucy Deconv (RLD) in Fourier light-field microscopy[1]. H is the forward projection matrix (PSF of Fourier light-field microscopy). $H^{-1}$ is the backward projection matrix derived from H. ii, View-To-Volume: A 2D U-Net based network (VCD[2]) maps the stacked views into volume in a self-supervised manner with the wave-optics model consistency. iii, Volume-To-Volume: Similar to the “volume refinement” strategy[3], an initial volume was first derived from once back projection operation in Deconv and then was refined by 3D U-Net with physics model consistency. iv, View-To-Volume (3D volumes-supervised): This approach adopts the same network structure in (**ii**) except that the loss function is computed from predicted volume and targe volume. v, LFINR: LFINR adopts a full-connected neural network to represents 3D volume under the supervision of input views. INR-based network queries spatial coordinates to generate intensity information at each position. **b,** Lateral and axial section planes in a 3D zebrafish embryo heart (cardiomyocyte nuclei labelled with GFP) reconstructed by RLD, 2D U-Net, 3D U-Net, 2D U-Net (with 3D volume priors) and LFINR. The Deconv results suffered from structural distortion and axial elongation; “2D U-Net” improved the lateral structural fidelity but yielded axial discontinuity; “3D U-Net” preserve the axial continuity with deconvolved 3D stack but failed to refine this volume, characterized by the axial elongation in the marked white rectangular area; “2D U-Net (with 3D volume priors)” reconstructed volume without artifacts and discontinuity with volumetric priors but exhibited blurry in lateral projections; LFINR performed realistic in lateral projection while mitigated the elongation and preserved high continuity in XZ plane. **More details seen in Supplementary Note 1.**

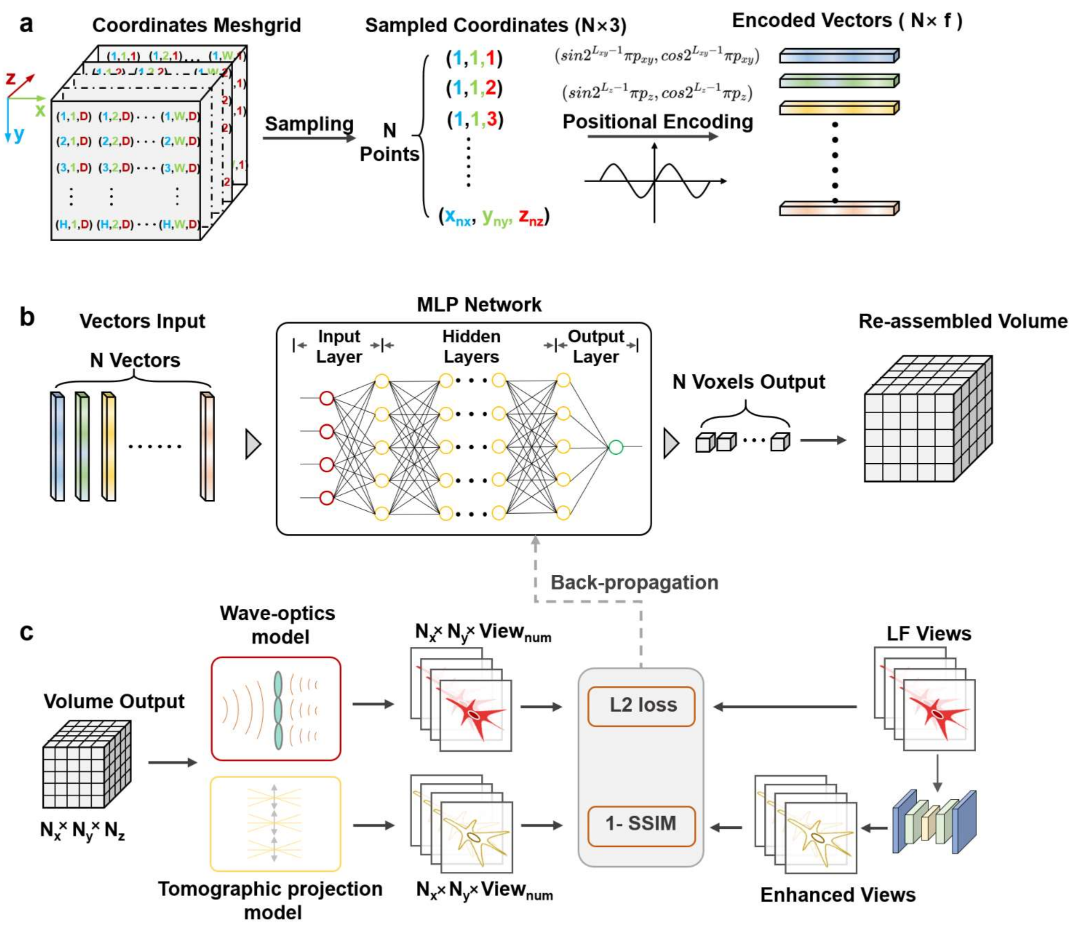


**Supplementary Figure 2**. **Data-flow in network optimization process**. **a,** Positional encoding on sampled coordinates. The sampled coordinates ($N$×3, where $N = N_x \times N_y \times N_z$) were encoded via Fourier positional encoding formula[4], producing encoded vectors with a size of ($N$×$f$). $f$ is channel number of the encoded feature and is equal to $3 + 3 \times 2 \times (p_x + p_y + p_z)$. **b,** MLP network inference. MLP network converted the input $N$ vectors to $N$ voxels. Then, reposition these voxels via their own spatial coordinates. MLP consists of input layer, hidden layers and output layer. The channel number of hidden layers was listed in **Supplementary Table 1**. The channel of output layer is 1 because of the grayscale images. **c,** Loss computation. The volume output with a size of $(N_x, N_y, N_z)$ was converted to dual projections with wave-optics model and tomographic projection model. The input LF views were also enhanced by pre-trained network. The dual projection and the two LF views were used to compute the hybrid loss function. Then, conduct back-propagation to update the parameters of MLP.

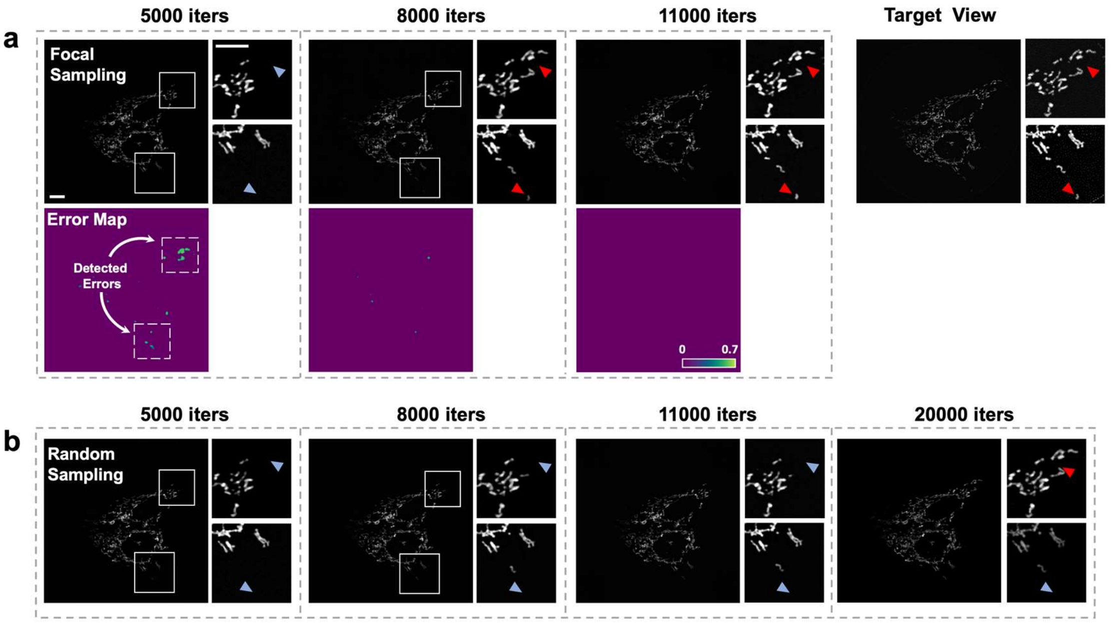


**Supplementary Figure 3**. **Dynamics sampling strategy significantly accelerate the convergence of network optimization**. **a**, The xy maximum intensity projections (MIPs) of mitochondria's LFINR reconstructions at different iterations (top row), along with their dynamic error maps compared to the target view (bottom row). The error maps were continuously updated to detect areas of missing signal, as highlighted by the white dotted boxes. This focal sampling strategy facilitated the rapid recovery of the missed weak signal from 5000 to 8000 iterations, as indicated by the changes from blue arrowheads to red ones. The LFINR projection has perfectly matched the ground truth without errors at the 11000$^{th}$ iterations. **b**, The xy MIPs of reconstructions by random sampling strategy. Without the spatial awareness on structural errors, conventional INR reconstruction based on random sampling strategy failed to recover the weak signals (indicated by the blue arrowheads), even after 20000$^{th}$ iterations. Scale bar: 10μm.

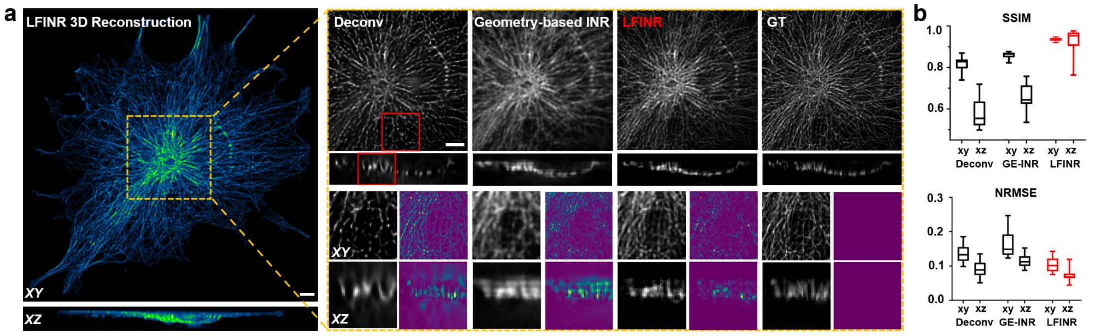


**Supplementary Figure 4**. **Comparative performances of Deconv, Geometry-model-based INR and LFINR on the reconstruction of simulated FLFM projection of microtubule**. **a,** The reconstruction results offered by FLFM Deconv (Deconv), geometry-based INR and LFINR. Geometry-model approach conducted the geometry projection in hybrid model rendering branch while LFINR adopted dual-projection rendering strategy, in network optimization process. **b,** The statistical results of SSIM and NRMSE of different approaches. GE-INR: geometry-model based approach. The mean SSIM are 0.818/0.579 (Deconv at XY/XZ plane), 0.859/0.655 (GE-INR at XY/XZ plane) and 0.936/0.927 (LFINR at XY/XZ plane). n > 5. The mean NRMSE are 0.136/0.091 (Deconv at XY/XZ plane), 0.165/0.115 (GE-INR at XY/XZ plane) and 0.105/0.072 (LFINR at XY/XZ plane). n > 5. Scale bars: 5 μm in (**a**, **b**).

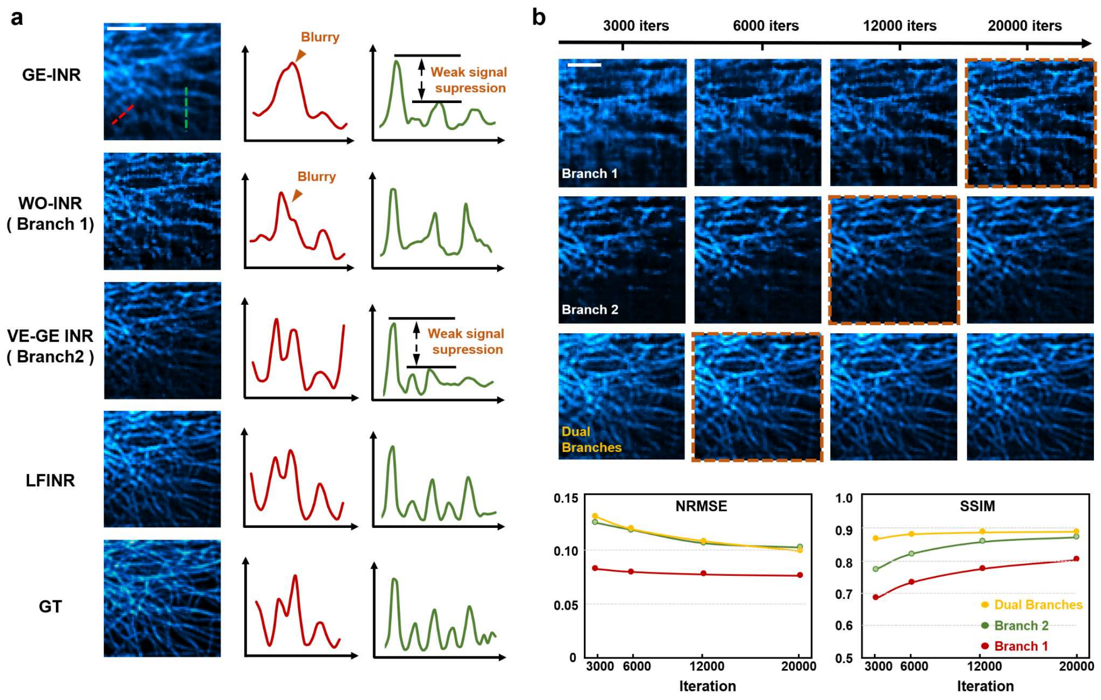


**Supplementary Figure 5. Performance evaluation of INR approaches with different rendering models. a,** Maximum intensity projections (MIPs) of microtubules reconstructed using geometry model with FLFM views ("GE-INR"), wave-optics based INR with FLFM views ("WO-INR", branch 1), geometry model with enhanced views ("VE-GE INR", branch 2), and LFINR (dual branches). The line intensity profiles (red and green lines) of the reconstructed signals from different rendering methods are compared, demonstrating the superior fidelity of LFINR with the hybrid model. **b,** The reconstruction results obtained using different rendering methods and their corresponding quality metrics (SSIM and NRMSE) across different network iteration stages. The brown dashed box indicated the earlier convergence in dual branches results. Scale bars: 10 μm in (**a**, **b**). More details are described in **Supplementary Note 3.**

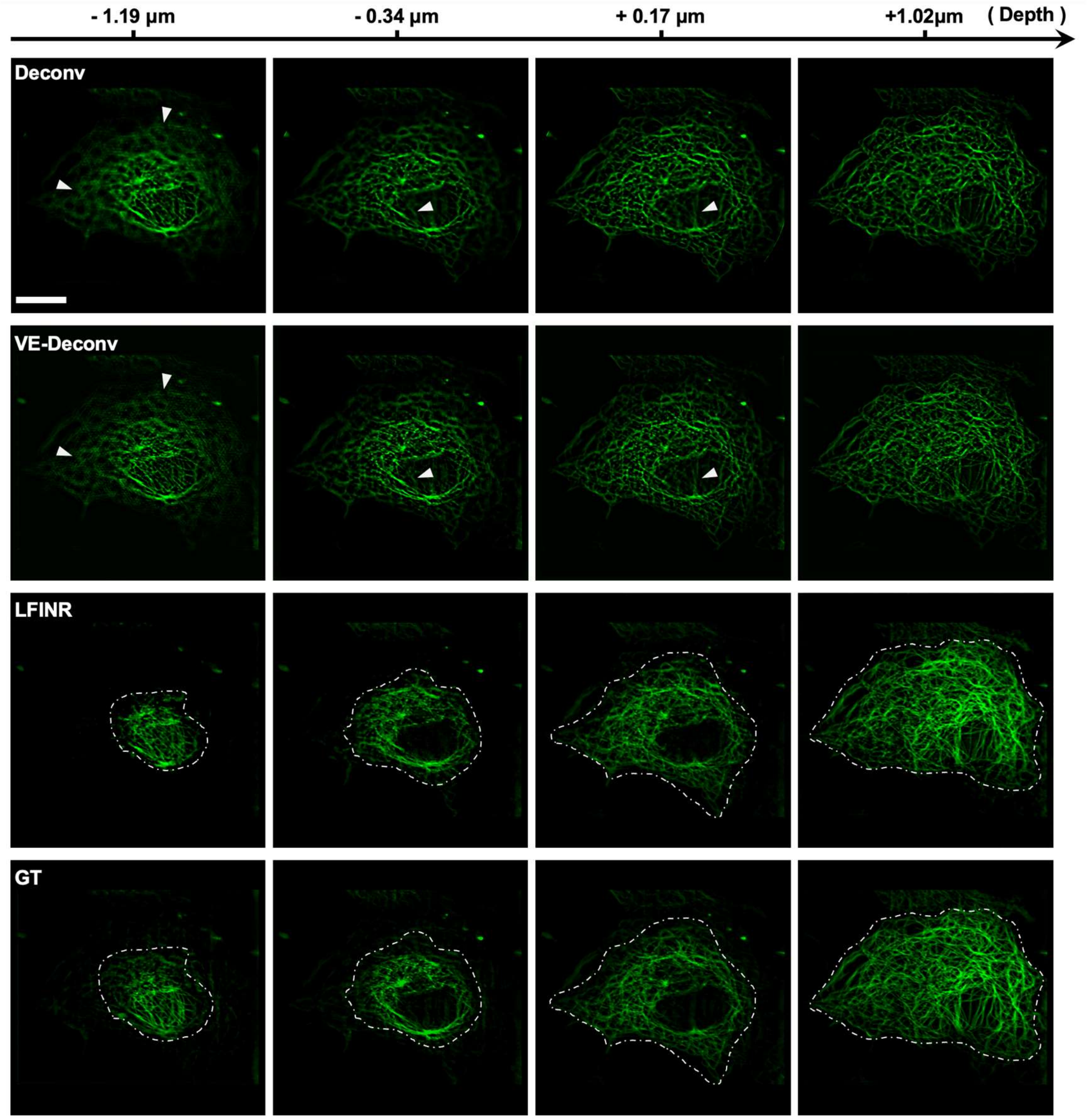


**Supplementary Figure 6**. **Comparison of the optical sectioning capability of deconvolution-based approaches and LFINR on microtubules data.** Deconv: FLFM Deconv. VE-Deconv: view-enhanced Deconv. The white arrowheads indicate the axial elongation of signals caused by Deconv approaches, resulting in compromised optical sectioning capability. Scale bar: 20 µm.

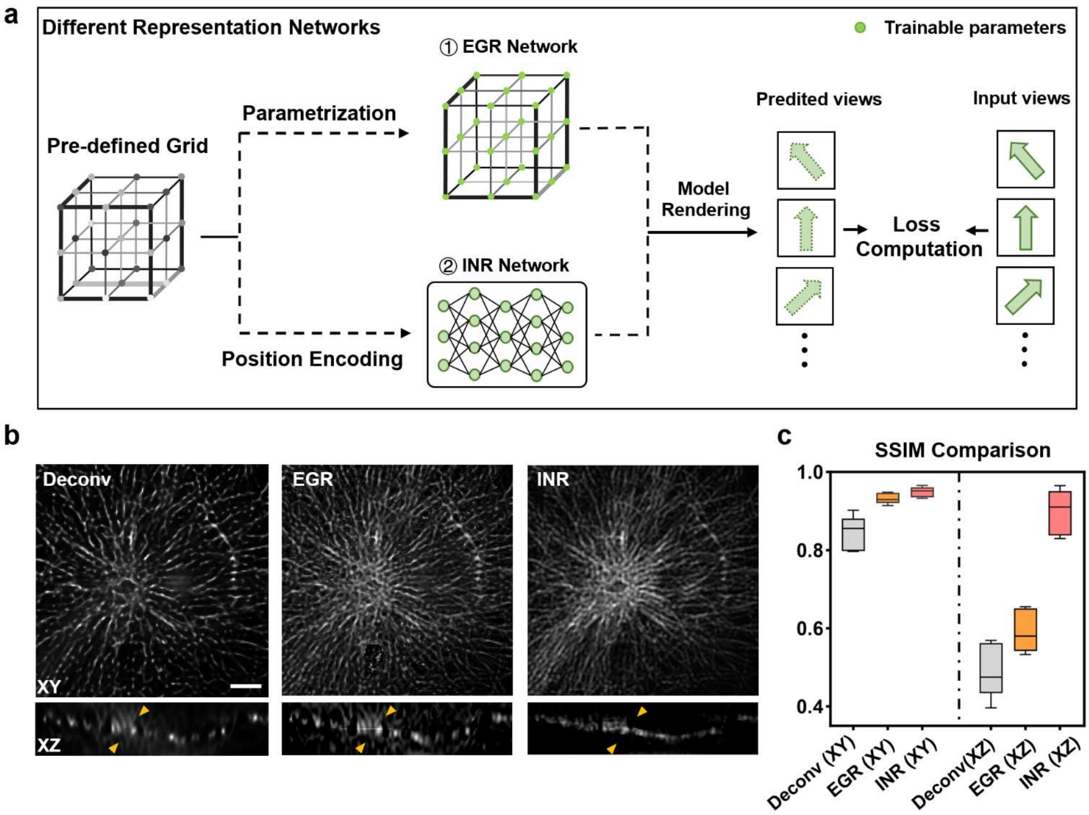


**Supplementary Figure 7. The axial enhancement offered by the continuity of implicit neural representation. a**, The framework of explicit grid-based representation (EGR) and implicit neural representation (INR). Similar with Deconv, EGR initialized a volume whose voxels can be optimized via loss function. While INR adopted a Multi-Layer Perceptron (MLP) to learn a function which converted the vectors from encoded coordinates to corresponding voxel values. **b**, The xy/xz projections of 3D microtubule reconstrued by FLFM Deconv (Deconv), EGR approach and INR approach. The intrinsic continuity of MLP facilitate to suppress the axial elongation and artifacts annotated with orange arrowheads. Scale bar: 5 µm. **c,** The SSIM statistical results of Deconv, EGR and INR reconstruction. The mean SSIM are 0.842/0.487 (Deconv at XY/XZ plane), 0.933/0.590 (EGR at XY/XZ plane) and 0.949/0.901 (INR at XY/XZ plane). $n > 5$.

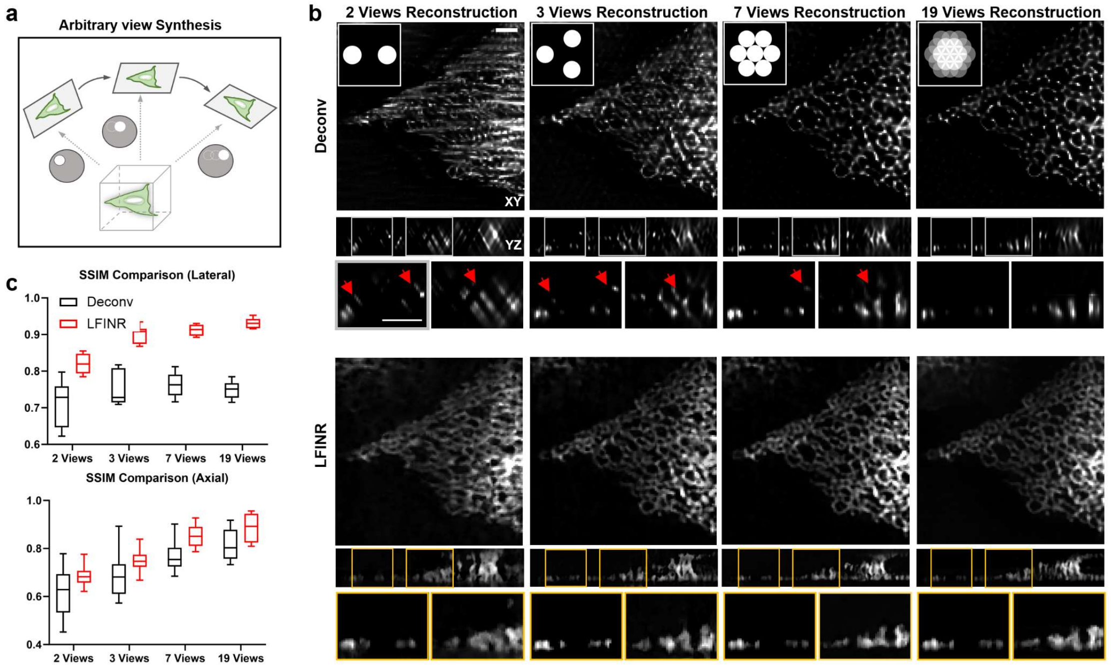


**Supplementary Figure 8. LFINR enables artifact suppression in 3D reconstructions with various angular sampling rates**. **a,** The schematic of generating multi-views with arbitrary position. To investigate the performance of LFINR in various angular sampling rates with the same spatial resolution, we adopted "sequential-projection" strategy to generate any number of FLFM view within the limited optical aperture. **b,** Reconstruction comparison via Deconv and LFINR under different numbers of FLFM views of endoplasmic reticulum (ER). Rows indicate the lateral projection/axial slice of Deconv results and LFINR prediction. Columns indicate the 3D reconstruction under different views. Deconv results exhibit directional distortion and unnatural details denoted by low SSIM value in **(c)**. LFINR results show centralized distribution of frequency components and improved SSIM (2 views: *xy*: 0.727 /0.818, *xz*: 0.629/0.680; 3 views: *xy*: 0.729 /0.906, *xz*: 0.682/0.746; 7 views: *xy*: 0.763 /0.915, *xz*: 0.757/0.853; 19 views: *xy*: 0.751 /0.929, *xz*: 0.800/0.890). Scale bar 3 μm in (**b**). The sample size of statistics results in **(c)** is more than 5.

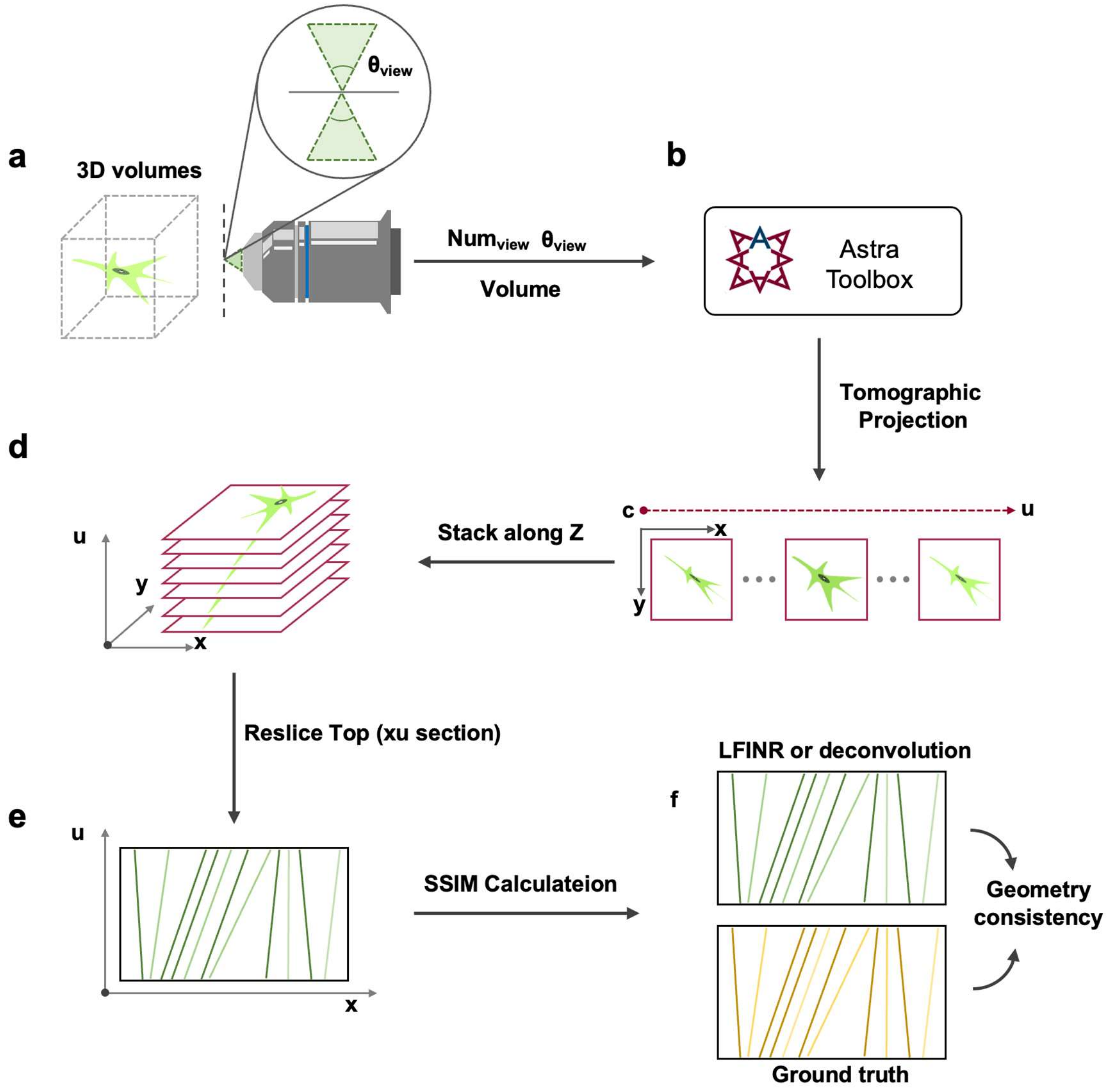


**Supplementary Figure 9. Workflow of geometry consistency computation.** To evaluate the validity of synthesized view offered by different approaches, we conducted structure similarity index measurement (SSIM) computation on generated epi-polar plane images (EPIs), following 6 steps: **a,** Obtain the view-range decided by objective' NA and input the 3D volumes (LFINR reconstruction, deconvolution-based results and ground truth stack). To ensure the angular continuity of generated EPIs, the number of views was set to 19 in this paper. **b**, Feed the volumes and hardware settings into Astra-toolbox[5] (a toolbox for 2D and 3D tomography). **c,** Conduct tomographic projection to generate 19 views with horizontal arrangement. **d**, Stack these views along z-axis. **e**. "Reslice" the stack and get *xu* section plane. **f**, Compute the SSIM metrics between LFINR or deconvolution-based EPIs and ground truth ones.

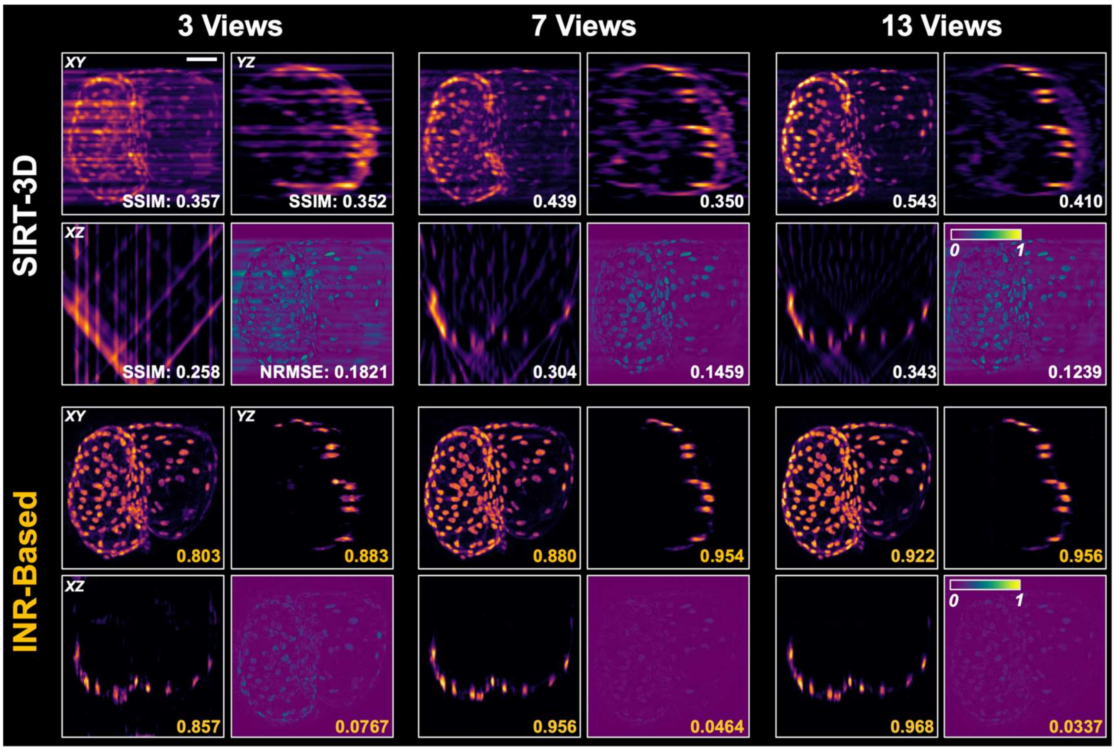


**Supplementary Figure 10**. **INR-based optical projection tomography (OPT) reconstruction with few angles**. 3D OPT reconstructions of zebrafish heart (labelled with nuclei) by SIRT-3D (top) and INR (bottom) approaches. Left to right: The *xy*/*xz*/*yz* planes of 3D reconstructions under different number of views. The similarity metric SSIM and error metric NRMSE are calculated at the lower right corner of images to indicate the significantly higher fidelity achieved with the INR-based approach compared to classic SIRT-3D. Scale bar: 30 μm**.** More details about data preparation and network settings can be found at **Supplementary Note 5.**

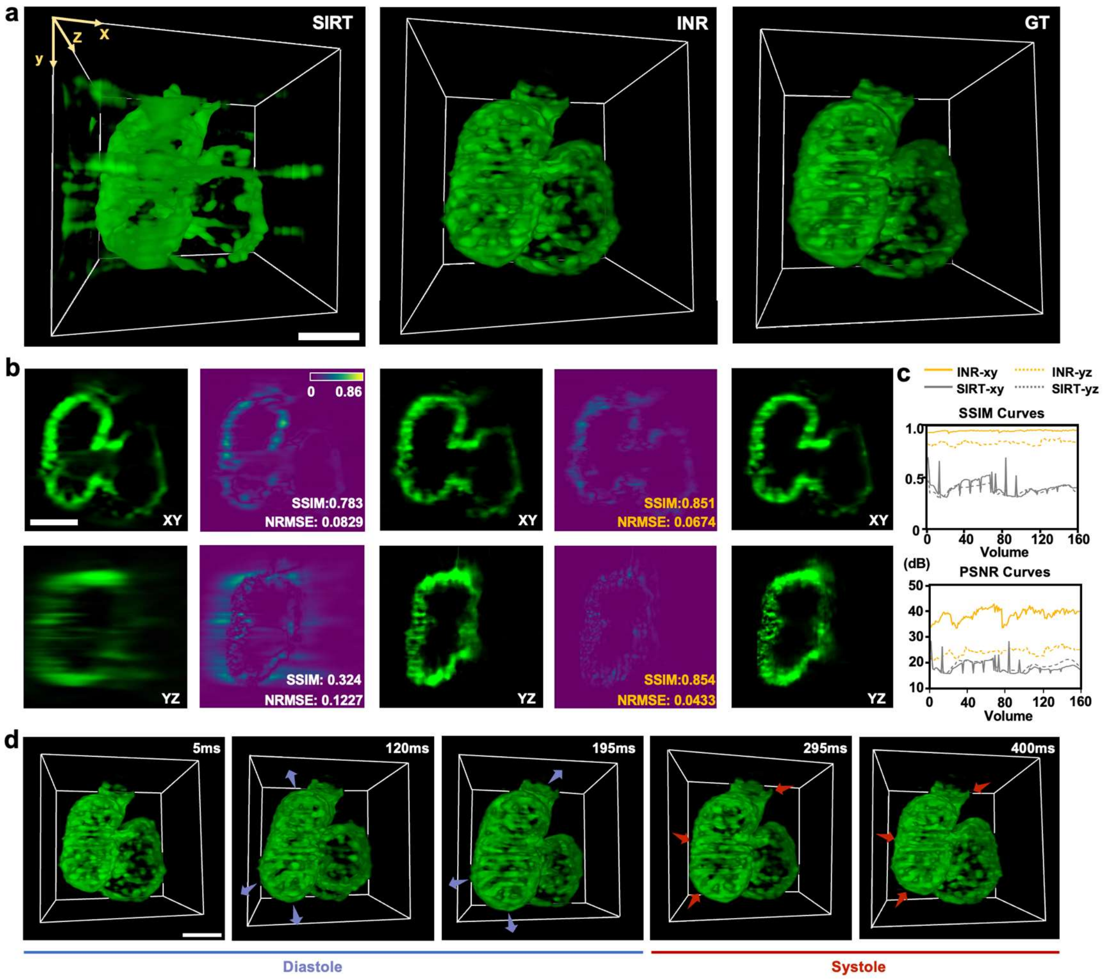


**Supplementary Figure 11. Comparison of the reconstruction of beating heart in a live zebrafish larva using conventional SIRT and meta-learning based INR**. **a,** Volume renderings of beating zebrafish heart (*Tg(cmlc2:gfp)*, 4 days post fertilization) reconstructed by SIRT-3D and INR approaches. The results are compared with ground truth (GT) data which are acquired by light-sheet fluorescence microscopy combined with retrospective gating algorithm[6]. **b,** Comparison of the orthogonal views and the corresponding error maps of the volume renderings shown in a. **c,** Comparison of reconstruction fidelity across 160 timepoints spanning two cardiac cycles was quantified by calculating SSIM and PSNR metrics using the GT as reference. **d,** The time sequence of the beating myocardium during diastole and systole in one cardiac cycle. Scale bar: 20 μm. More details about data preparation and network configurations can be found at **Supplementary Note 5**.

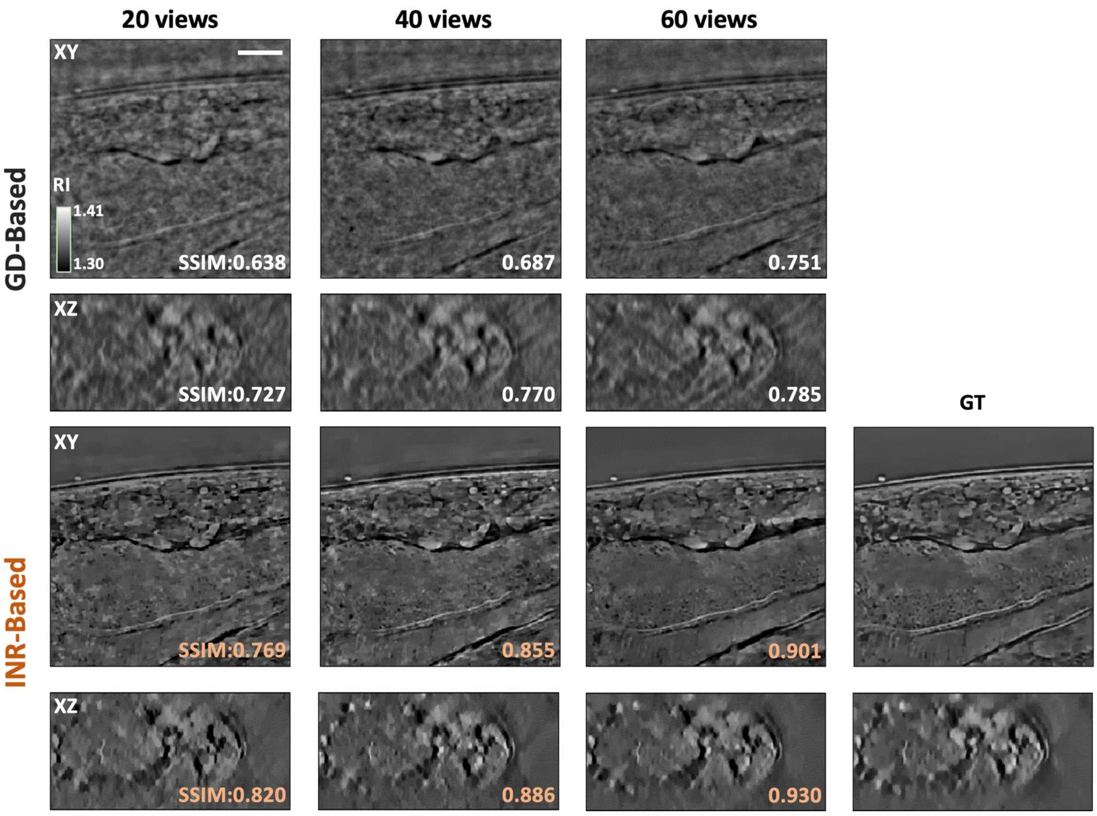


**Supplementary Figure 12**. **INR-based Fourier ptychographic microscopy (FPM) reconstruction with suppressed artifacts.** FPM 3D reconstructions by gradient-decent (GD, top) and INR (bottom) approaches relied on Multi-Layer Born[7] approximation on *C. elegans*. Left to right: the xy/xz planes of FPM results under different number of illumination sources. Scale bar: 10 µm. More details about data preparation and network configurations can be found at **Supplementary Note 5.**

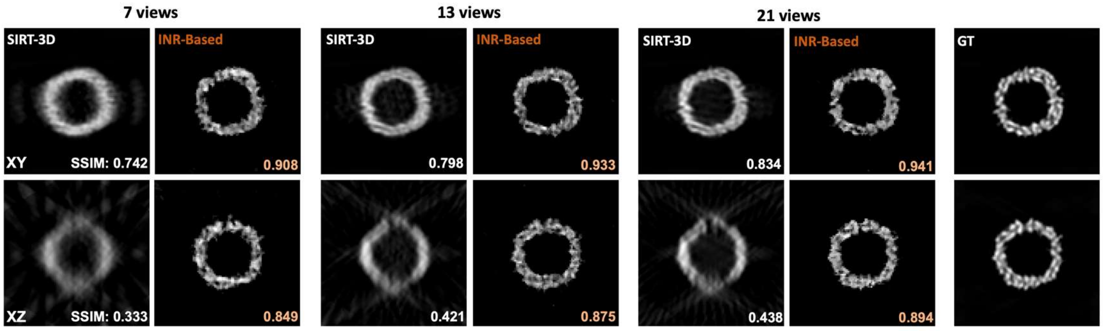


**Supplementary Figure 13**. **INR-based cryo-electron tomography (Cryo-ET) reconstruction.** The tomographic 3D reconstruction of apoferritin by SIRT-3D and our INR approach. Left to right: the xy/xz planes of the 3D results reconstructed with 7, 13, and 21 tilting angles. INR Cryo-ET showed significantly better reconstruction quality under all the view settings. More details about data preparation and network configurations can be found at **Supplementary Note 5.**

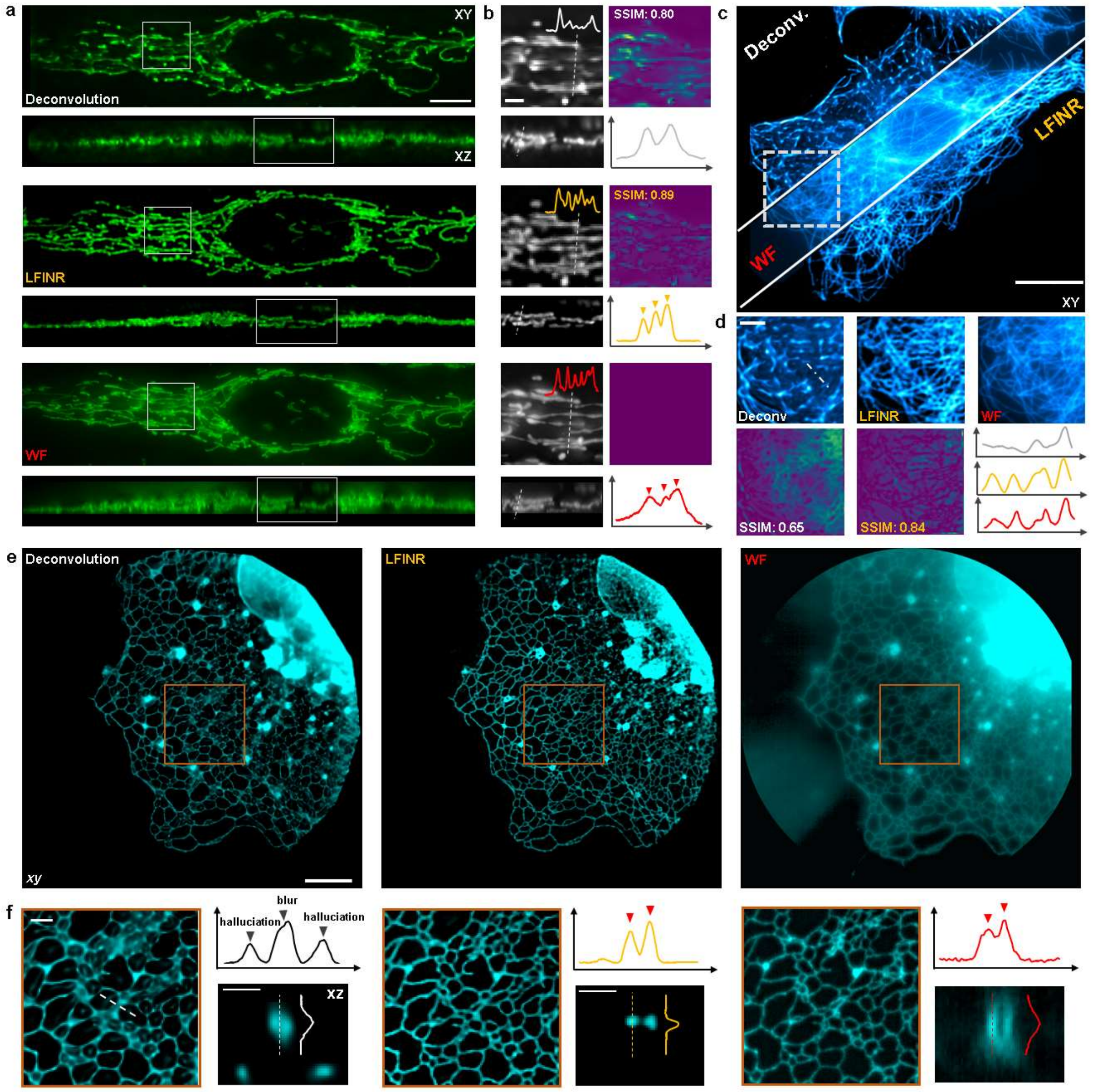


**Supplementary Figure 14. The reconstruction fidelity of LFINR on diverse structures. a,** The results of mitochondria (tagged with Cox4-EGFP) via Deconv, LFINR and wide-field (WF) microscope. **b,** The magnified view of white rectangular regions shown in **(a).** The error maps and intensity profiles along dotted lines are shown. **c,** The xy-MIP of microtubule (tagged with anti-tubulin-alexa488) in fixed U2OS cells. **d,** The higher-magnification views of the region marked with white dashed-box in **(b)**. **e**, The ER (tagged with Sec61β-EGFP) network in a Cos7 cell, by Deconv, LFINR and WF microscope. **f,** Magnified high-resolution views from the brown box in **(e).** The gray arrowheads indicated the quality degradation of Deconv results while the red arrowheads showed the high similarity between LFINR results and WF acquisition. Scale bars: 10 μm in (**a**), 2 μm in (**b**), 20 μm in (**c**), 5 μm in (**d**), 10 μm in (**e**), 2 μm in (**f**).

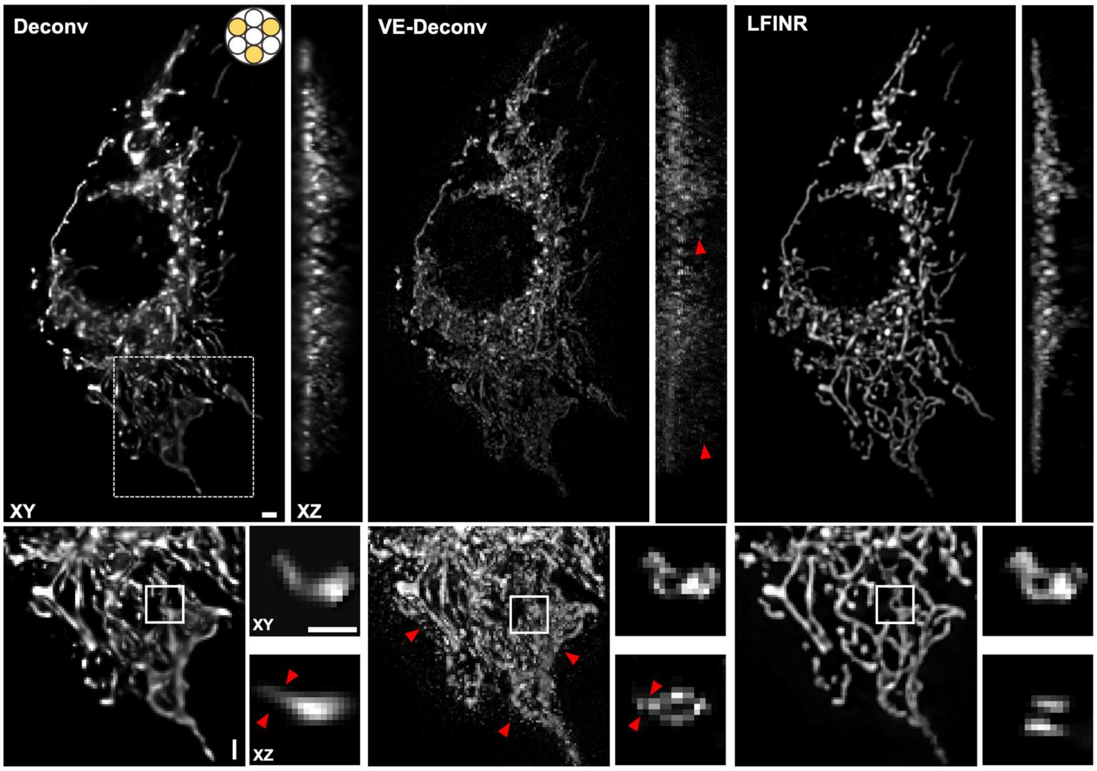


**Supplementary Figure 15**. **The non-full aperture FLFM reconstruction of fixed mitochondria matrix using Deconv, VE-Deconv and LFINR.** The red arrowheads indicated the striped reconstruction artifacts in Deconv and VE-Deconv when using only three input views. Scale bars: 2 μm.

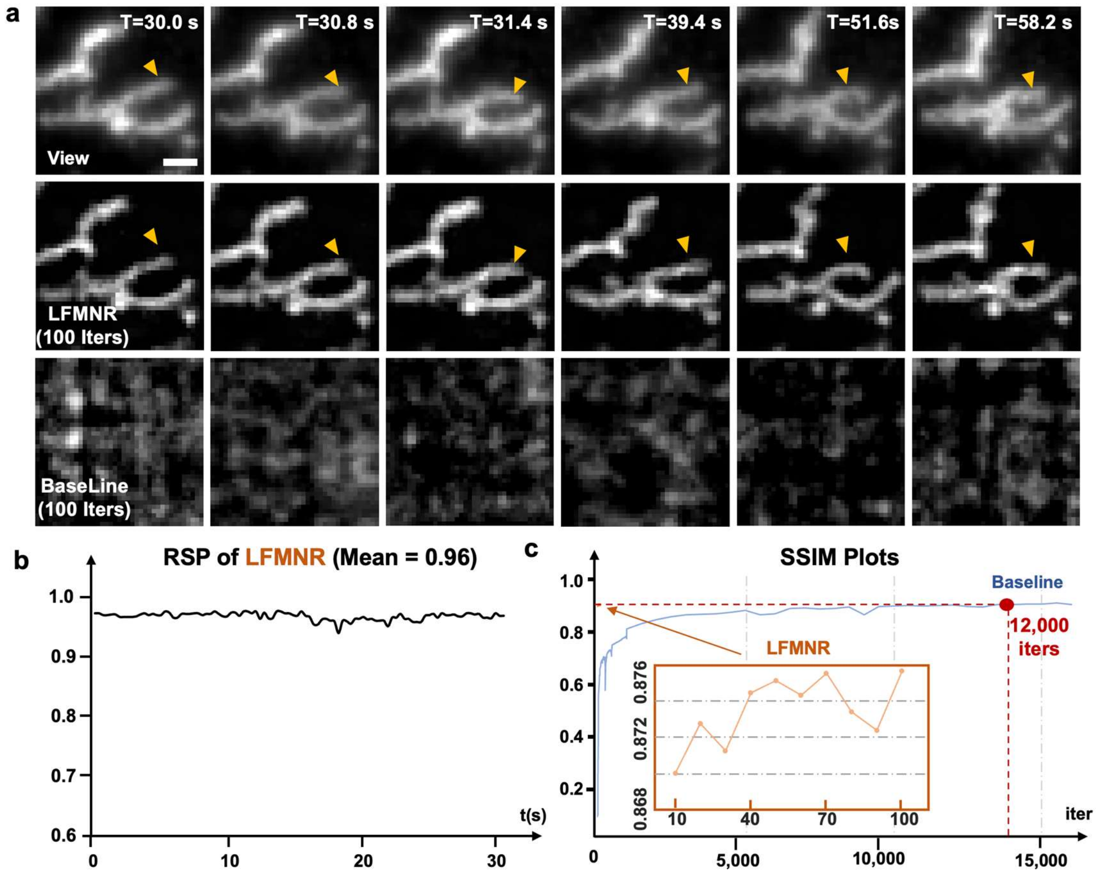


**Supplementary Figure 16**. **Light-field neural representation with meta-learning (LFMNR) accelerates network optimization in 4D (3D space + 1D time) imaging.** **a**, Ablation study on meta-learning. Top row: FLFM views of mitochondria. Center row: The xy MIPs of LFMNR results. Bottom row: The xy MIPs of baseline INR method without meta-learning. The orange arrowheads indicated that the local wiggling of mitochondria was successfully reconstructed by LFMNR after 100 iterations. In contrast, the baseline approach failed to reconstruct any effective signals, owing to the slow convergence. Scale bar: 2 μm. **b**, The RSP[8] metrics calculated between LFMNR and LFMNR view, validating the high structure fidelity of LFMNR for reconstructing mitochondrial wiggling shown in (**a**). **c**, The structural similarity index measure (SSIM) plots of LFMNR and baseline during network optimization. LFMNR achieved fast convergence within 100 iterations while baseline required ~12000 iterations for achieving similar SSIM.

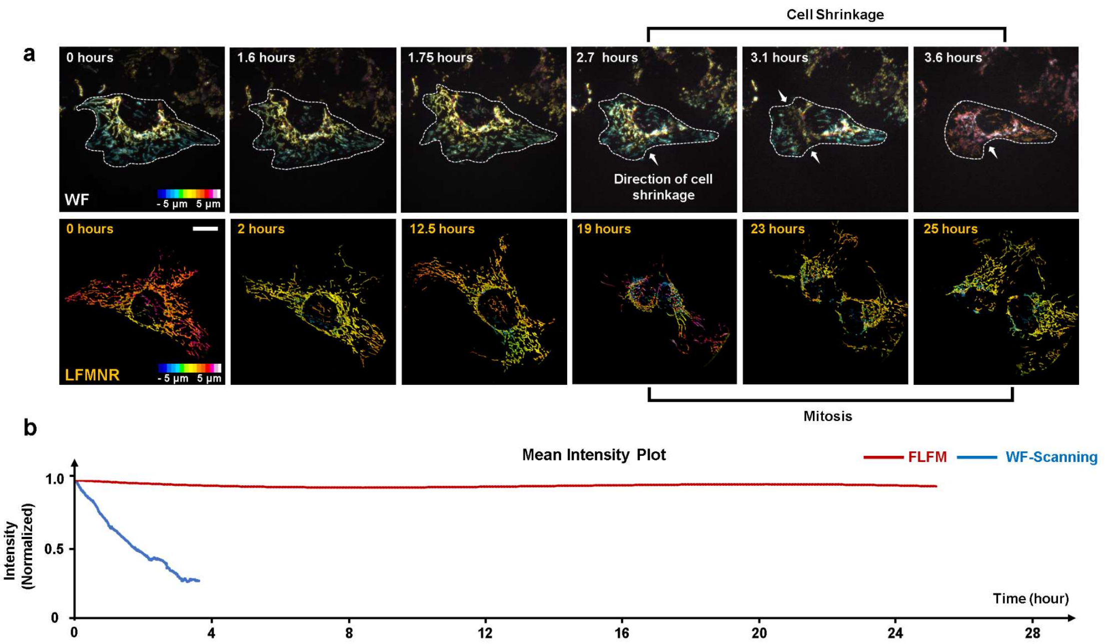


**Supplementary Figure 17. Long-term imaging of mitochondria dynamics in live U2OS cells.** Comparison of phototoxicity and photobleaching between 3D wide-field imaging and LFMNR imaging. The 3D WF and LF imaging were performed under the same light dose per frame (7.1mJ/cm$^2$) and volumetric imaging time interval (15 s). To acquire a volume, light-field imaging captures only a single frame (~250 ms total), while wide-field imaging requires axial scanning with 51 frames (~15 s total). **a,** The MIPs of mitochondria in live U2OS cells (depth encoded by color) are shown to compare the cell states at different time points. The dotted lines mark the cell boundaries in WF results and the arrows indicate the direction of cell shrinkage due to the phototoxicity. The cell imaged using LFMNR divided after 19 hours of observation, with a total recording duration of 25 hours capturing 6000 volumes, validating the ultra-low phototoxicity of LFMNR. Scale bar: 20 μm. **b,** Comparisons of the photobleaching rates between WF scanning approach and LFMNR.

### Supplementary Note 1: Discussion about classical model and deep-learning based light-field reconstruction algorithms

Fourier light field microscopy (FLFM) enables rapid volumetric imaging of dynamic biological processes through capturing multiple views of the dynamic signals in one quick snapshot. Recovering the 3D signals from the 2D view measurements is crucial to FLFM. However, this process is ill-posed due to information loss caused by depth and angular-wise encoding, as well as the sum projection in the FLFM forward model, similar to sparse-view and limited-angle tomographic imaging[1, 9, 10].

Deconv ("*Deconv*", **Supplementary Figure 1a**) addresses the problem merely relying on the prior of the physics model (point-spread function of the FLFM system), thereby being naturally influenced by the limitations of limited angles and sparse views. For example, Richard-Lucy Deconv[1, 11] suffered from the artifacts and axial elongation when solving such inverse problem as shown in **Supplementary Figures 1b, 4a** and **7b**. Volume-supervised approaches circumvent the limitations imposed by hardware settings through introducing external data priors on spatial structures or sample distributions. However, they encountered challenges in model generalization on various sample[12], and required access to numerous high-quality paired 2D-3D data which were laborious and required complex imaging system[13]. This requirement of 3D data acquisition would be hard for fast biological dynamics. Furthermore, the mapping process in 3D-supervised networks[13, 14] lacks interpretability, hindering the seamless integration of deep learning networks with physical models.

Self-supervised learning-based approaches offer an alternative solution to this problem. To our best knowledge, most self-supervised algorithms incorporate the physics model with a learnable neural network model under the constraint of physics consistency[15]. However, there is a lack of discussion about self-supervised implementation and the corresponding mechanism in solving the inverse problem of FLFM 3D reconstruction. Here, we conducted two experiments to investigate this potential principle:

***First, "2D to 3D" mapping function.*** A practical way to obtain a self-supervised 3D reconstruction network by following the paradigm[16, 17] , which involves building the mapping function (CNN, convolution neural network) from source domain (2D views, in this study) to target domain (3D volume, in this study) with the constrains of physics model consistency. For instance, as shown in (ii) of **Supplementary Figure 1a**, a 2D U-Net network[2] (VCD) widely used in light-field 3D reconstruction was constructed to convert the stacked 2D views to 3D volume. During training process, the loss function can be computed from input FLFM views and re-projected views obtained from the convolution of network prediction and FLFM PSF. To build this network, we generated approximately 1500 stacked views with a size of 240×240×19 from 3D stacks of cardiomyocyte nuclei with the 19-view hardware settings. The learning rate was set to $5\times10^{-4}$, and the loss function was the mean square error (MSE) between input views and network projection. The results (referred to as "*ii. 2D U-Net*" in **Supplementary Figure 1b**) showed higher structural fidelity in lateral projections but severe discontinuity in axial sections. We attributed the lateral enhancement to the CNN's fitting ability under the lateral constraints of each input 2D view, while the axial discontinuity originated from the lack of volumetric priors. This network, serving as the mapping function from "2D to 3D", consists of numerous 2D filters that first convert stacked views into high-dimensional features and then reshape these extracted features with the size of 3D volume. During this process, there is no explicit or implicit volumetric information or representation, and no continuity constraints are applied across different channels of the extracted features. Therefore, this algorithm design, while effective in enhancing lateral features, requires extensive volumetric priors to achieve a natural transformation from "view" to "depth".

***Second, "3D to 3D" refinement.*** Another way to achieve self-supervised learning involved conducting a back-projection operation to convert the encoded observations into an "initial guess" of target domain and refining the guess results by a network with the constrains of physics consistency[18]. Here, following the paradigm in multi-view 3D reconstruction[3], the process begins by using the transposed point spread function (PSF) to convert input views into a coarse volume. Subsequently, a 3D network (3D U-Net, **Supplementary Note Table 1**) refines this volume with the constraint of minimizing the loss computation between network projections and input views ("*iii. Volume-To-Volume*", **Supplementary Figure 1**). We adopted Adam optimizer with a learning rate of $5\times10^{-4}$. The training data contained ~1500 FLFM patches. As the results shown in "*iii. 2D U-Net*" of **Supplementary Figure 1b**, the axial discontinuity was removed with the aid of initial "guessed-volume" derived from single back projection. However, the axial elongation and artifacts occurred, denoting such refined network still suffered the defects in solving limited-angle & sparse-view 3D reconstruction. Such "*Volume-To-Volume*" process contains non-linear network mapping and forward projection loss computing, which is equal to refine the initial guess volume in image domain using the loss computed from input views and forward projections. This iterative refinement process is highly similar to Deconv approach, whose volume was refined iteratively with the guidance of discrepancy between projected views. Besides, considering the low-frequency bias of network[19], such self-supervised refined algorithm tended to block the high frequency information, resulting in a the blurry effect in rectangular area of "*iii*, **Supplementary Figure 1b**", which was similar to images denoising results in untrained network[20]. Consequently, without additional data priors, this image-refining strategy also exhibited certain limitations.

Based on the observed limitations in the aforementioned experiments, we conclude that the primary challenge in self-supervised 3D reconstruction lies in preserving the continuity of reconstructed signals while mitigating axial elongation and artifacts inherent in solving inverse problems.

Back to the essence of 3D reconstruction in FLFM, the ultimate goal is to obtain 3D signal distribution from 2D multi-view image. While the discussed approaches aimed to learn the potential mapping function and reconstruct volumes indirectly, an alternative direct approach was to represent the high-dimensional signal with a continuous neural function[4, 21-24], termed implicit neural representation (INR). This continuity in learned function derived from the continuous coordinates input and the continuous fitting ability on complex representation function of Multi-Layer Perceptron (MLP) [25-27] , which could generate continuous views with geometry consistency and act an "implicit" regularization on signal distribution. Inspired by this, we developed LFINR that represent the 3D complex biological signals with the multi-view observation captured by high-speed Fourier light-field microscopy. As **Supplementary Figure 1a.v** shows, the MLP served as the underlying volumetric function and optimized with the physics model consistency. Compared to relying solely on the physics model prior in Deconv or self-supervised mapping learning strategy, LFINR integrated the continuity bias served as a "model prior" to regularize the potential solution space, and constrain the signal distribution in implicit domain. Additionally, the novel view synthesis ability brought by the implicit neural representation was highly correlated in solving multi-view 3D inverse problem. Therefore, LFINR ("*v. LFINR*", **Supplementary Figure 1b**) achieved artifact-free 3D reconstruction with reduced axial elongation and enhanced axial continuity similar to the volume supervised approach.

| **Alias** | **Layer** | **Operations** |
|---|---|---|
| Input | InputLayer | Convert to ViewStack |
| U-Net (Encoder) | EN_1 | conv3d(n8 k3 s1 p1)->Relu->conv3d(n8 k3 s1 p1) |
| | | DownSample |
| | EN_2 | conv3d(n16 k3 s1 p1)->Relu->conv3d(n16 k3 s1 p1) |
| | | DownSample |
| | | conv3d(n32 k3 s1 p1)->Relu->conv3d(n16 k3 s1 p1) |
| U-Net (Middle) | Middle | UpSample |
| | | concat(Middle,EN_2) |
| U-Net (Decoder) | | conv3d(n16 k3 s1 p1)->Relu->conv3d(n16 k3 s1 p1) |
| | DE_1 | UpSample |
| | | concat(DE_1,EN_1) |
| | DE_2 | conv3d(n16 k3 s1 p1)->Relu->conv3d(n16 k3 s1 p1) |
| Output | output | conv3d(n1 k3 s1 p1)->Relu |

**Supplementary Note Table 1. Network structure of U-Net 3D**

**Supplementary Note 2：Self-supervised view-enhancement network**

The FLFM views suffered resolution degradation due to blur modulation induced by point spread function (PSF) distortion and depth-wise defocus degradation. Such degradation would further decrease the SBR (signal-to-background ratio) and influence the reconstruction quality by classical model-based approach[28, 29]. Taking this into consideration, we conducted 2D-Deconv operation to remove the blur in FLFM views (as shown in "*For static scene (pre-Deconv)*" **Supplementary Note Figure 1**), which produced high-contrast FLFM views. However, for numerous FLFM images in capturing fast dynamics, such pre-Deconv approaches would influence the throughput of data-processing. Therefore, we built a view-enhancement network to fast deconvolved sequential FLFM views. The detailed steps as follow:

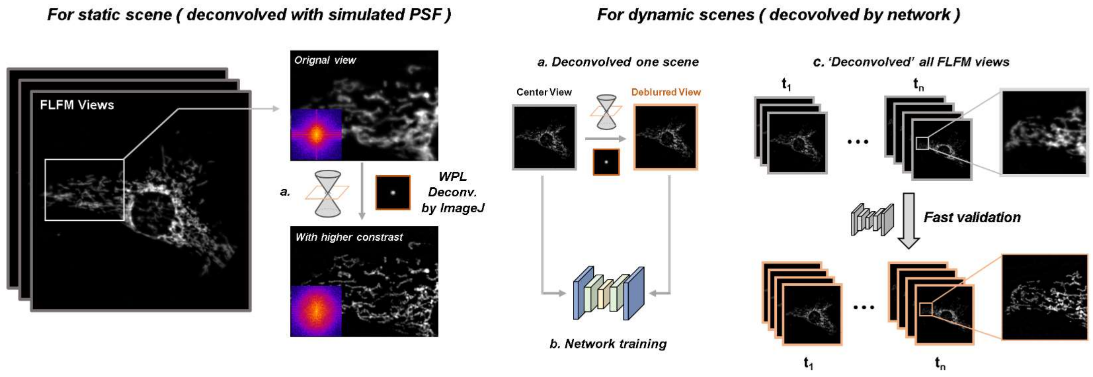


**Supplementary Note Figure 1. The generation process of enhanced FLFM views.** For static scene, the simulated PSF was used to remove the blur in FLFM views, which enhanced the structural contrast and expanded the effective frequency bandwidth of the views in the Fourier domain. For dynamic scenes, a network was used to accelerate the deblurring operations.

*For static scene:*

***a***. Using PSF generator in *Fiji* to obtain 2D PSF which is equivalent to the sub-aperture PSF of center view in FLFM.

***b***. Applied 2D Deconv algorithm ("*Parallel iterative Deconv*" in *Fiji*) on various views to remove the blur with the simulated 2D PSF. Such Deconv operation could remove the blur in input views and produce higher contrast.

*For dynamic scene*

***a***. Deconvolved the one scene of FLFM captures follow the steps in processing static scene.

***b***. Crop the paired views (original views and deconvolved views) and train a U-Net for view-enhancement.

***c***. Deconvolved sequential images with trained network.

The architecture of view-enhancement network was illustrated in **Supplementary Note Figure 2**. In this study, we employed Adam as the optimizer with a learning rate of $4\times10^{-4}$. The number of training pairs of one scene was usually 256~512 (varied with different samples) and the lateral size was 96×96. The batch size was set to 16. Total time of network training was 4~8 minutes.

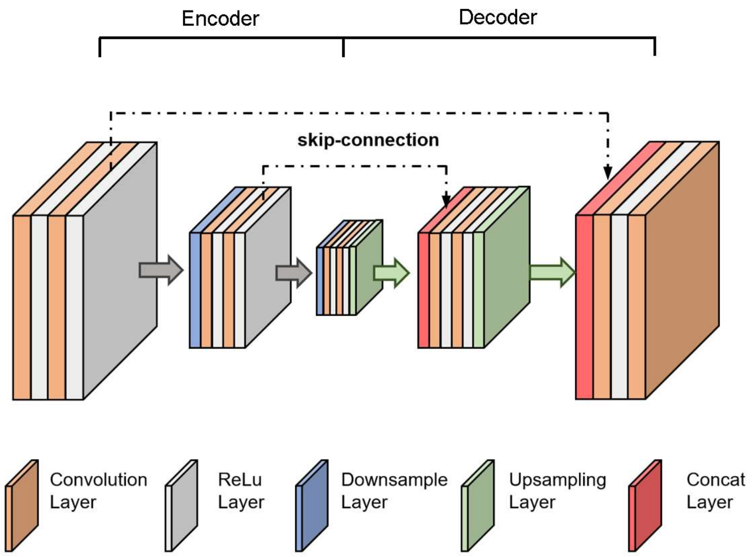


**Supplementary Note Figure 2. Network structure of view-enhancement network.** We adopted U-Net composed of encoder and decoder to deblur the input views. Each encoder block contains 2 convolution layers with parameters n (channel number) and f (size of convolution kernel) while each decoder block contains a bicubic upsampling layer and 2 convolution layers. The filter size and basic channel number in our experiment were set to 3 and 32, respectively.

### Supplementary Note 3：The mechanism of hybrid physics rendering model

Light-field 3D reconstruction is an inverse problem whose potential solution space is varied with different optical setups but can be constrained by the prior knowledge. FLFM Deconv approach failed to yield high-quality reconstruction only under the constrain from FLFM PSF (point-spread function), manifested as the axial elongation and artifacts in limited-angle reconstruction (**Fig. 2**). So, to remove such axial degradation in multi-view reconstruction problem, more constrains need to be obtained.

The first step is to obtain the prior from the continuity bias, namely, the model prior of implicit neural representation (INR) network. In INR, MLP (multilayer-perceptron) acting as a function approximator can fit the complex representation function in a continuous manner[24, 26, 27]. With this prior, INR could produce the predicted views with high geometry consistency, which prompted the quality of 3D scene representation in multi-view reconstruction. In the FLFM, one plausible way for INR application is to replace the ray-casting model in NeRF with PSF. The results in **Supplementary Figure 5** show that wave-optics model based INR (WO-INR) mitigate the blur and signal loss in geometry model based INR (GE-INR). This improvement derived from the more accurate computation model in WO-INR. However, with enough iterations, WO-INR still shows poor resolution. This poor quality of WO-INR is derived from the less constrains in solving 3D reconstruction problem.

Except of the prior knowledge, the complexity of forward model also deicides the difficulty in solving inverse problem. Though geometry model based algorithm ignores the blur in captured views, but this model can provide faster reconstruction speed[9] due to the less complexity and is prevalent in natural scene representation[4]. This simplified model could yield promising results when no blur effect in views (**Supplementary Figure 10**). This inspired us to remove the blur in FLFM captured views to mitigate the mismatching between FLFM and geometry projective model ("VE-GR INR" in in **Supplementary Figure 5**). The results provided better spatial resolution compared with WO-INR. Besides, the simplified model reduced the iteration times to achieve better results. However, although the blur effects were mitigated, the processed views were not perfect to enable recovering the accurate 3D signal distribution because of the incompatibility between FLFM imaging and tomographic projective model, denoted by the weak-signal missing.

Based on above analysis, we proposed physics-based hybrid rendering model that introduces the "wave-geometry" model constraints. Combining with the continuity bias of INR, our method further tightens the possible solutions space of 3D reconstruction and achieved faster convergence (**Supplementary Figure 5**). Specifically, the better matching degree between FLFM views and wave-optics model describe the intensity transmission in forward process, which mitigated the signal depression occurred in VE-GE INR. While the blur effects and artifacts in WO-INR results are removed by deblurred views with higher structural contrast and the pixel-wise line projection model. The mutual promotion in dual branches achieves a synergistic effect where the whole is greater than the sum of its parts.

**Supplementary Note 4: Self-supervised denoising network.**

Due to the inherent noise and fluorescent intensity decay presented in living cell imaging, we employed a self-supervised denoising network to preprocess the raw light field (LF) views.

For raw LF sequence with high frame rate, for example, the fast voltage imaging (100 Hz), we adopted Noise2Noise[30] strategy to remove the noise, whose mechanism is based on the mean gradient of massive paired noisy data is approximately equal to the true gradient. With the time-redundancy in LF sequence, the training paired data could be obtained by interlacedly sampling on the whole time-sequence. In this study, the raw captures firstly split into two sequences. Then, we conducted a fixed interval sampling ($\Delta t$=10) along the sequences to generate training pairs.

For other application with lower frame rate, for example, observing the cellular apoptosis process, the spatial redundancy[31] was utilized as a prior to remove the noise. Specifically, the captured sequences were first uniformly sampled at fixed time intervals. Subsequently, the extracted views from these sampled LF sequences were down-sampled to generate paired training data using a neighbor subsampler. In this study, pixels from the upper left and bottom right corners within each local 2×2 area were selected to create such training pairs.

After the data preparation, we used a light-weight U-Net to remove the noise. The network followed the structure of view-enhanced network (**Supplementary Note Fig. 2**). We used Adam optimizer and mean square error (MSE) as loss function. The model was built with Keras and trained on RTX 3090. The training typically takes 5~10 minutes.

**Supplementary Note 5: INR for various limited-angle tomographic reconstruction tasks.**

Our proposed INR-based reconstruction approach can inference the 3D volume from multi 2D projections with limited & sparse views. In addition to solving the ill-posed 3D reconstruction problem in light-field microscopy, INR can be further extended to a variety of other 3D computational imaging modalities. Here, we implemented INR-based OPT, FPM and Cryo-ET reconstructions, to fully validate INR's superior versality as well as high performance.

First, we demonstrated the utility of proposed method in limited-angle OPT. OPT obtains the multi-view projections of observed sample by sequentially rotating sample or optical imaging path[32, 33]. Generally, in order to improve the volumetric imaging speed and reduce the photo phototoxicity, the angle-range and rotating times are simultaneously restricted. In this study, we used high-resolution 3D volume data (cardiomyocyte's nuclei of zebrafish labeled with *Tg* (*myl7:nls-gfp*), captured by confocal imaging system with a ×20/0.75W objective and beating zebrafish heart (*Tg*(*cmlc2:gfp*), 4 days post fertilization) capture by light-sheet fluorescence microscopy combined with retrospective gating algorithm[6]) to generate multi-views within limited angular range (-40°~+40°). The rotation axis was set to y-axis. Then, we used SIRT-3D[5], a widely used traditional algorithm, and our INR-based algorithm to reconstruction 3D volume from these simulated OPT views. As shown in **Supplementary Figure 10**, INR yields artifacts-free reconstruction even under 3 input views and produces finer structural details in axial planes, while SIRT-3D suffered from the striped artifacts and axial blur caused by sparse angular sampling and missing cone in frequency domain. As shown in **Supplementary Figure 11,** INR-based approach achieved better structure fidelity across cardiac cycles compared with SIRT-3D**.** The network architecture follows the paradigm of LFINR but differs in forward projection model and network hyper parameters: In OPT forward projection, we first calculated angle-wise affine transformation matrixes, and then conducted sum-projection along detection direction. The MLP contains 6 fully-connected (FC) layers with 128 channels. The order of position encoding was set to 8 for lateral plane and 6 for $z$ axis.

We also evaluated the performance of INR for reconstructing FPM and Cryo-ET data. In FPM, multi-angle projection views of a sample could be obtained by sequential illumination from multiple angles, therefore, reducing the number of images collected would benefit the acceleration of volumetric imaging speed, thereby improving the system's temporal throughput. To demonstrate the reconstruction capability of INR for strongly scattering samples under limited illumination angles, we utilized the *C. elegans* volume data[34] to generate multiple views. With the settings of a 1.05 NA objective and 0.9 NA maximum illumination angle, we compared the reconstructed 3D refractive index (RI) of the traditional gradient-decent based algorithm (GD) and INR based algorithm under 20, 40, 60 views respectively (**Supplementary Figure 12**), both algorithms relied on the Multi-Layer Born (MLB) approximation[7] as the forward projection model. During reconstruction, the network's output was constrained between 1.33 and 1.52. It could be seen that INR-based method could produce results with fewer artifacts and much higher contrast, and it can still obtain finer structural details under just 20 views, while the traditional algorithm suffered serious artifacts and axial blur due to the sparse angle illumination. Apart from the different forward projection model used, the network architecture remains similar to LFINR, comprising a MLP with 5 FC layers, including a skip connection at the 3rd layer, and 16 channels.

As for Cryo-ET, the multi-view projections were obtained by tilting the sample sequentially. During this process, the allowable electron dose was distributed to each projection, which indicated that reducing the number of acquired tilts would contribute to preventing structure damage and enhancing SNR[35]. Here, to evaluate the performance of INR in solving sparse-view reconstruction, we used the apoferritin volume data[36] to generate multiple tilting images. It should be noted that we preferred to discuss the generalizability of INR in solving sparse view reconstruction in cryo-ET. A comprehensive and exhaustive evaluation on cryo-ET reconstruction is beyond the scope of this work. Therefore, we adopted a simplified forward model of cryo- ET by tiling the particle structure along y-axis without the consideration of the impacts of noise and low contrast in real cryo-ET images (These problems can be solved with image pre-processing, like images matching & averaging, CTF modulation.). The tilting range here follows the conventional settings, -60°~+60°. We used SIRT-3D and INR method to reconstruct 3D volume from 7, 13 and 21 tilts (**Supplementary Figure 13**). As the results show, the continuity bias of INR offered the artifacts suppression and axial resolution enhancement compared with blurry results of SIRT-3D. The number of FC layers of MLP was set to 6 and the number of channels was 64. The maximum order of position encoding for $xy$ and $z$ were both 6.

**Supplementary Table 1: Network hyperparameters of LFINR**

| | 3D Grid Size | Positional Encoding Order | Channel Number | Number of Layer | Weights-coefficients in loss function |
|---|---|---|---|---|---|
| Microtubules | 499×499×41 | $L_{xy} = L_z = 8$ | 128 | 8 | $\lambda_1$=0.1<br>$\lambda_2$=0.03<br>$\lambda_3$=0.03 |
| Endoplasmic reticulum | 499×499×41 | $L_{xy} = L_z = 8$ | 128 | 8 | $\lambda_1$=0.1<br>$\lambda_2$=0.01<br>$\lambda_3$=0.05 |
| Mitochondria matrix | 499×499×51 | $L_{xy} = L_z = 8$ | 128 | 4 | $\lambda_1$=0.1<br>$\lambda_2$=0.01<br>$\lambda_3$=0.1 |
| Cardiomyocyte nuclei | 307×307×51 | $L_{xy} = L_z = 8$ | 128 | 8 | $\lambda_1$=0.1<br>$\lambda_2$=0<br>$\lambda_3$=0 |
| Lysosome | 307×307×51 | $L_{xy} = L_z = 8$ | 128 | 4 | $\lambda_1$=0.1<br>$\lambda_2$=0.02<br>$\lambda_3$=0.1 |
| Terminal bulb | 215×215×91 | $L_{xy} = 8$<br>$L_z = 6$ | 128 | 8 | $\lambda_1$=0.1<br>$\lambda_2$=0.001<br>$\lambda_3$=0.01 |
| Pharyngeal muscle | 319×319×91 | $L_{xy} = L_z = 6$ | 128 | 6 | $\lambda_1$=0.1<br>$\lambda_2$=0.001<br>$\lambda_3$=0.1 |
| Zebrafish brain | 380×308×101 | $L_{xy} = L_z = 8$ | 128 | 8 | $\lambda_1$=0.1<br>$\lambda_2$=0.001<br>$\lambda_3$=0.01 |

**Supplementary Table 2：Optical parameters of Fourier light-field microscope**

| Alias | Figures | Objective | Fourier Lens | Microlens Array | FOV[1] (Field of View) | Theoretical Resolution[1] |
|---|---|---|---|---|---|---|
| 3-views Setup | Fig. 2d | 100×/1.4 | *f*: 300 mm | *Pitch*: 3.25 mm,<br>*f*: 120 mm | 81.25 µm | $R_{xy}$: 0.485 µm<br>$R_z$:0.775 µm |
| | Fig. 3h | 100×/1.4 | *f*: 250 mm | *Pitch*: 3.25 mm,<br>*f*: 120 mm | 67.71 µm | Rxy: 0.404 µm<br>$R_z$:0.538 µm |
| 7-views Setup | Fig. 2b<br>Figs. 3a,3d<br>Figs. 4i, j, m | 60×/1.5 | *f*: 250 mm | *Pitch*: 3.25 mm,<br>*f*: 120 mm | 112.85 µm | $R_{xy}$:0.673 µm<br>$R_z$: 0.863 µm |
| | Fig. 5e<br>pharyngeal<br>muscle | 30×/1.05 | *f*: 250 mm | *Pitch*: 3.63mm,<br>*f*: 120 mm | 252.08 µm | $R_{xy}$: 1.205 µm<br>$R_z$: 2.766 µm |
| 19-views Setup | Fig. 5e<br>terminal<br>bulb | 30×/1.05 | *f*: 250 mm | *Pitch*: 3.25 mm,<br>*f*: 120 mm | 225.69 µm | $R_{xy}$: 1.538 µm<br>$R_z$: 1.972 µm |
| | Fig. 2g | 20×/0.5 | *f*: 250 mm | *Pitch*: 2.0 mm,<br>*f*: 150 mm | 166.67 µm | $R_{xy}$: 3.281 µm<br>$R_z$: 10.253 µm |
| 27-views Setup[37] | Figs. 5a, b | 25×/1.05 | *f*:160 mm | *Pitch*: 1.3 mm,<br>*f*: 26 mm | 800 µm | $R_{xy}$: 3.4 µm<br>$R_z$: 5 µm |
